# Probing for Intentions: The Early Readiness Potential Does Not Reflect Awareness of Motor Preparation

**DOI:** 10.1101/2023.08.10.552835

**Authors:** Jake Gavenas, Aaron Schurger, Uri Maoz

## Abstract

Voluntary actions are typically preceded by the Readiness Potential (RP), a negative midfrontal EEG deflection that begins ∼2 seconds before movement. What cognitive and neural process the RP reflects and how it relates to conscious intention remain unclear due to conflicting findings. We investigated the neural basis and cognitive significance of the RP in a novel probe-based paradigm. Contrary to prior reports, we found that pre-probe RP buildups were not related to reported awareness of motor preparation. Computational modeling suggested that the best explanation for these results is via metacognitive access to stochastic accumulation. Reported preparation was also related to beta desynchronization over contralateral motor cortex shortly before probe onset. We conclude that the RP may be metacognitively accessible in response to external task demands but does not reflect the onset of a conscious intention. We discuss implications of these findings for voluntary action initiation and intention awareness.

**Highlights:**

1. We investigate the mechanisms underlying voluntary action initiation in a new probe paradigm.
2. Contrary to prior results, the readiness does not reflect reported awareness of motor preparation.
3. Computational modeling supports stochastic accumulation over linear ballistic accumulation and classic RP models.
4. Reported awareness of motor preparation may emerge from metacognitive access to stochastic accumulation.
5. Time-frequency analysis suggests reported awareness may also relate to pre-probe beta desynchronization.

## 1 Introduction

Voluntary actions are typically preceded by a slow buildup of aggregate neural activity in midfrontal regions^1–5^. Single-unit activity in the supplementary motor area (SMA) and anterior cingulate cortex that ramped up or down in advance of movement onset was also demonstrated before voluntary action^6^. The most well-studied of these slow-ramping signals is the readiness potential (RP), an event-related potential that consists of a negative deflection in electroencephalography (EEG) that begins around 2 seconds before self-initiated movements^1,7^. Despite the decades that have passed since the RP’s discovery, the kind of process that this signal reflects, and its cognitive significance and role in action initiation remain contested.

Traditionally, the onset of the RP was thought to reflect a discrete neuro-cognitive event such as the beginning of motor preparation^1^, or the onset of an unconscious decision to move^2,7,8^. In contrast, more recent work suggested that the RP may instead reflect an integration-to-bound process of autocorrelated noise, which then resembles a gradual deflection when aligned to threshold-crossing and back-averaged^9–14^ (see^15^ for review). The so-called Stochastic Accumulator model thus parsimoniously accounts for the early rise and for the shape of the RP as well as for the distribution of wait times in unconstrained self-initiated action. It has further made novel predictions that were empirically confirmed^9^, and we recently demonstrated that a spiking neural network extension of stochastic accumulation can parsimoniously explain pre-movement activity at the single-neuro and network levels^16^. However, Bogler and colleagues^17^ suggested that evidence specific for stochastic accumulation is still lacking, because other models can account for many of the same phenomena as stochastic accumulation. For instance, they note that linear ballistic accumulation—a process in which a monotonic linear buildup with a random slope triggers movement upon crossing a threshold^18^—can also account for the RP’s shape and waiting-time distributions. Unlike the traditional model of the RP, the onset of ramping in stochastic accumulation and linear ballistic-accumulation models does not reflect a specific event that initiates the ramping. The three models entail different implications for the neural bases of spontaneous voluntary action, the implementation of conscious control, and the utility of the RP and similar signals for movement prediction such as for brain-computer interfaces. Distinguishing between them is therefore of paramount importance.

Efforts to establish what kind of process underlies the RP are further complicated by conflicting results concerning the RP’s cognitive significance^19^. Studies of self-initiated action often have participants report the onset of the conscious urge or intention to move (referred to as W-Time) using the position of a rotating clock^2,5,20^. Though several studies have found that the onset of the RP does not correlate with W-Time as reported using the clock method^5,21,22^, recent studies employing an alternate method of assessing participants’ subjective state found that the reported awareness of conscious intention may indeed be related to RP onset^23,24^ (though see^25^ for a contrary report). For instance, Parés-Pujolràs and colleagues^23^ utilized an online-reporting paradigm in which participants were randomly probed before they moved^26,27^ and asked whether they had already begun preparing to move. Using this method, Parés-Pujolràs and colleagues found RP-like deflections specifically before probes when the participants reported having begun preparing to move. Those deflections were absent when the participants did not report preparing to move. Similarly, Schultze-Kraft and colleagues^24^ used an online decoding algorithm based on the RP to deliver probes when the RP was present or absent. They found that reported awareness of preparation was more likely when their decoder indicated an RP was present compared to absent.

These results suggest that the RP may specifically relate to the conscious intention to move. Considering recent criticisms of the clock method^20,28–31^, it may be the case that probe methods, rather than clock methods, are the more appropriate avenue for investigating the onset of conscious intention. Nevertheless, it is unclear how these results influence the debate regarding what kind of process underlies the RP. Furthermore, probe methods inherently disrupt the movement-generation process, so they cannot determine when probes are delivered in relation to RP onset or to the onset of the movement that would have taken place had the probe not been delivered. It is therefore unclear whether the apparent link between the RP and reported awareness of intention apply to the early RP or only to its latest stages. Furthermore, decreases in beta power (desynchronization) in contralateral motor cortex also occur prior to self-initiated actions and appear to reflect motor preparation^7,32,33^. It was further claimed that beta desynchronization—rather than the RP—is related to awareness of motor preparation^25^, although evidence supporting that hypothesis remains tentative.

Here we developed a novel probe paradigm to investigate (1) whether the RP actually reflects the onset of conscious intention and (2) what neuronal processes the RP reflects. Participants were probed while they made self-paced movements and were instructed to inhibit their movement and report whether they were already preparing to move when the probe occurred. Importantly, our design enabled us to tease apart failures to inhibit—i.e., when motor commands were already ballistic and participants could no longer inhibit their movement—from earlier stages. If the early RP is related to conscious intentions, we would predict an RP-like buildup before probes on trials when participants successfully inhibit their movement and then report that they were indeed preparing to move. However, if pre-probe EEG amplitude does not co-vary with participants’ subjective reports, after controlling for failures-to-inhibit, it would suggest that the early RP does not relate to subjective awareness of motor preparation.

## 2 Methods

### 2.1 Participants

Twenty-four participants were recruited from the student population of Chapman University (Orange, CA) (mean age 19.5, range: 18-22, 71% female, 92% right-handed). Written informed consent was obtained from participants before they participated in the experiment. All methods were approved by the local ethics committee (Chapman University IRB-20-122).

### 2.2 Paradigm

We opted for a trial-based paradigm in line with computational models of self-initiated action. In every trial, Participants were instructed to wait approximately 3 s and then press spacebar whenever they wanted. Participants always used their right hands regardless of handedness. Importantly, they were further instructed to fully inhibit their movement if they heard an auditory tone (the probe; 0.1 seconds at 500 Hz; randomly timed via a shifted gamma distribution (4, 0.75) that began after 2 seconds; these parameters were selected to result in roughly 50% probe trials). There were therefore 5 types of possible outcomes in every trial (Fig. 1). (1) If participants moved before 2.5 s had elapsed from trial onset, they were warned that they moved too quickly. These were designated *Error Trials*. (2) If participants moved before probe onset, the experiment would automatically advance to the next trial. These were designated *Movement Trials*. (3) If the probe went off before participants moved, they were asked whether they were already preparing to move when the probe occurred. If they answered “yes,” these probe trials were designated *Prep Trials*. (4) If they answered “no,” these were designated *No Prep Trials*. Finally, (5) if the participants pressed the button within 200 ms after probe onset, the trial was marked as ‘failure to inhibit’ (FTI; see Data retention section below)).

**Figure 1:**
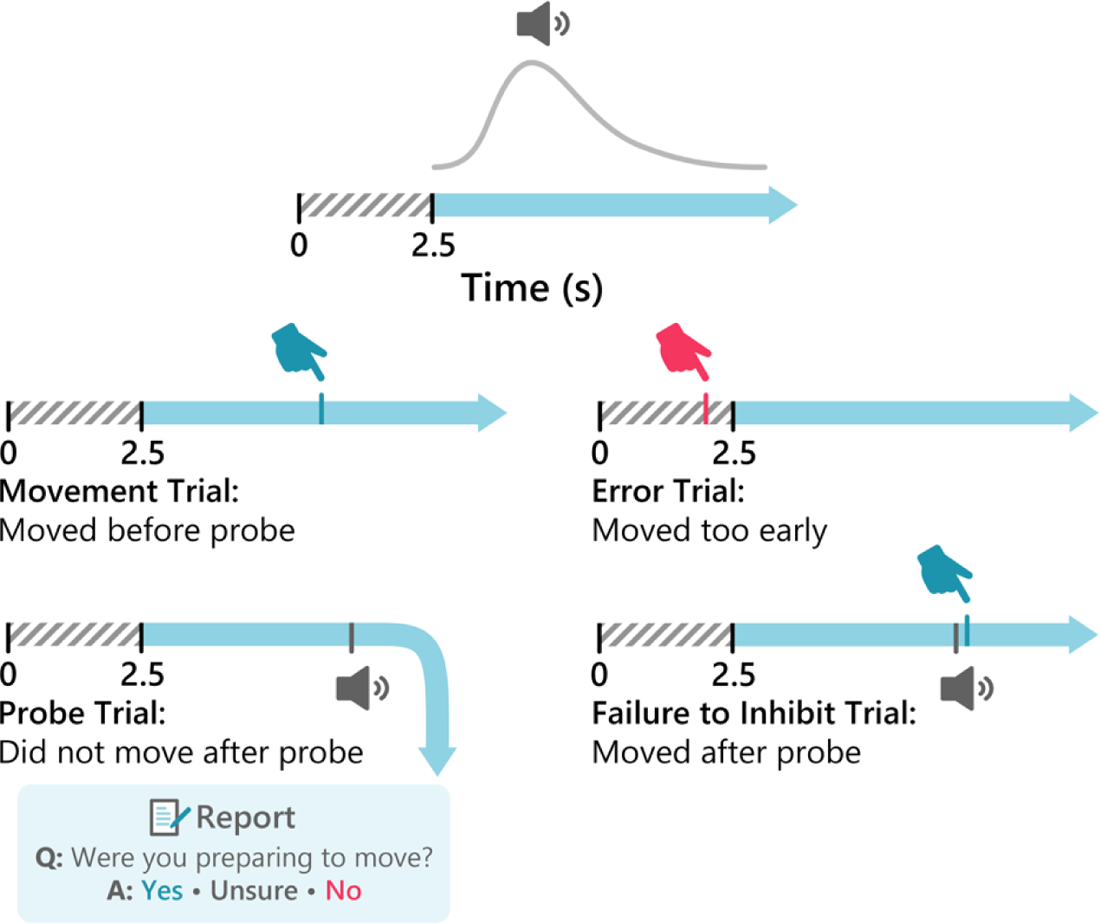
Paradigm. Participants were instructed to move whenever they felt like it after around 3 seconds had elapsed. At the beginning of each trial, the timing of an auditory probe was randomly generated according to a gamma distribution (top). Participants were instructed to inhibit movement following the probe and then indicate whether they were preparing to move at probe onset. These left four outcomes on each trial: movement trials, in which participants move before designated probe time; error trials, in which participants did not wait at least 2.5 seconds to move; probe trials, in which participants hear the probe and do not move following the probe; and failure-to-inhibit (FTI) trials, in which participants are unable to inhibit their movement following probe (presumably because the probe came too close to when they would have moved. Following the probe, participants were asked whether they were preparing to move. If they responded “yes”, the trial was designated a Preparation (Prep) trial, and if they responded “no,” the trial was designated a No Preparation (No Prep) trial. The rare trials where the participants responded “unsure” were omitted.

Regarding the movement, participants were instructed to press the spacebar with their right index finger spontaneously, without pre-planning, whenever they felt like it after waiting for around 3 seconds from the onset of the trial. However, they were further instructed not to count the seconds but to just estimate the time. For the first 40 trials, we let the participant get used to the paradigm, so we did not include probes. Hence, participants just carried out self-initiated action, while fixating on a white cross on a gray screen. After those initial trials, participants completed 20 practice trials of the main experiment (Fig. 1). After participants confirmed they understood the task instructions and required no more practice, we began the main experiment. We specifically instructed the participants not to let the possibility of a probe change how they generated actions throughout the experiment. In probe and FTI trials, 1 s after probe onset, participants were presented with a new screen and asked, “Were you preparing to move when the tone occurred?” Participants responded by pressing 1 (no), 2 (unsure), or 3 (yes) with their left hand. “Yes” responses were counted as “preparation” trials, and “No” responses were counted as “no preparation” trials. “Unsure” trials were not included in further analyses (see below). After practicing for 20 trials, participants were asked if they understood the experiment protocol, had the instructions repeated to them, and then they continued to the main experiment.

The actual experiment was broken into blocks of 20 trials each. And the participants were given a break between blocks. Our IRB approval set a limit of 2 h on our experimental session duration. Hence, after EEG setup and practice trials, we ran as many blocks as we could fit into the 2 h session. Participants completed between 8 and 10 blocks, and hence between 160 and 200 trials.

We were concerned that prior results linking the RP to access to consciousness were confounded by trials where participants were just about to move when the probe went off^23^ and thus could no longer inhibit their action. To account for such cases, we instructed participants to *fully inhibit all actions upon hearing the probe.* Hence, trials where the motor process had become ballistic would result in button presses after probe onset, which we termed failures-to-inhibit (FTI). Hence, in our paradigm, we could tease apart probe trials that included ballistic button presses (before which RPs would be expected) from probe trials in which participants’ responses were based on metacognitive access to the underlying neural process.

### 2.3 EEG recording and preprocessing

EEG was recorded online at 2048 Hz using a BioSemi 64-channel system and then read into MATLAB (MathWorks) R2019B and preprocessed using the FieldTrip software package. Data for each trial were epoched between 3.5 seconds before to 2 seconds after movement onset (movement trials) or tone onset (probe trials). First, all trials were visually inspected for artifacts, keeping the type of trial blind; trials containing artifacts were rejected (3.3±3.7% of all trials, range: 0-14%). We also removed electrodes that had poor signal-to-noise ratio based on visual inspection. Data were then downsampled to 200 Hz and bandpass filtered between 0.1 Hz and 35 Hz (filtered using Fieldtrip Defaults; Butterworth IIR filter). Using a 0.1 Hz high-pass filter has recently been suggested to reduce the impact of infra-slow oscillations (<0.1 Hz) when investigating the RP and other slow cortical potentials^34,35^. A non-filtered set of data was retained for time-frequency analysis. Independent Components Analysis (RunICA algorithm; FieldTrip implementation) was then used to visually remove artifacts corresponding to blinks and eye movements. The ICA procedure results in an unmixing matrix, which was then also used to remove those components from the unfiltered data. Finally, we re-referenced the EEG data to the overall mean of the remaining electrodes. For ERP analysis, we baseline-corrected the data by subtracting average EEG amplitude 2.5 to 2 seconds before movement, following prior studies^23^.

### 2.4 Time-frequency decomposition

Time-frequency analysis was conducted using custom scripts^36^ in MATLAB. We decomposed the data into the time-frequency domain using complex Morelet wavelets, ranging from 0.5 to 40 Hz. Wavelets started at 4 cycles and increased to 13 cycles as the frequency increased. Power was taken as the squared absolute value of the resulting complex time-series. For power in specific frequency bands, the frequencies that fell within that band (delta: 0.5-4 Hz; theta: 4-7 Hz; alpha: 7-12 Hz; low beta: 12-20 Hz; high beta: 20-30 Hz; gamma: 30-40 Hz) were averaged together.

### 2.5 Data retention

For EEG analyses, we omitted trials according to the following determined-in-advance criteria: (1) we removed probe trials on which the participants reported that they were ‘unsure’ whether they were preparing to move (3.6±3.5% of all trials, range: 0-11%), (2) we removed ‘error’ trials on which participants pressed the button less than 2.5 seconds after trial onset (3.4±2.9% of all trials, range: 0-11%), and (3) we removed ‘failure to inhibit’ trials on which participants pressed the button more than 200 milliseconds after probe onset (4.0±5.4% of all trials, range: 0-19%). 200 milliseconds before movement has been identified as the “point of no return” for self-initiated movements^37^. Therefore, movements following the probe by more than 200 milliseconds likely reflect lapses of attention rather than actual failures of inhibition. We used 200 ms as our cut-off to be conservative, even though the distribution of post-probe button presses in our analysis seemed to fall after about 250 milliseconds (see Fig. 2C). We retained the remaining data (besides those omitted from visualizing EEG; see EEG recording and preprocessing for details) for statistical analyses.

**Figure 2:**
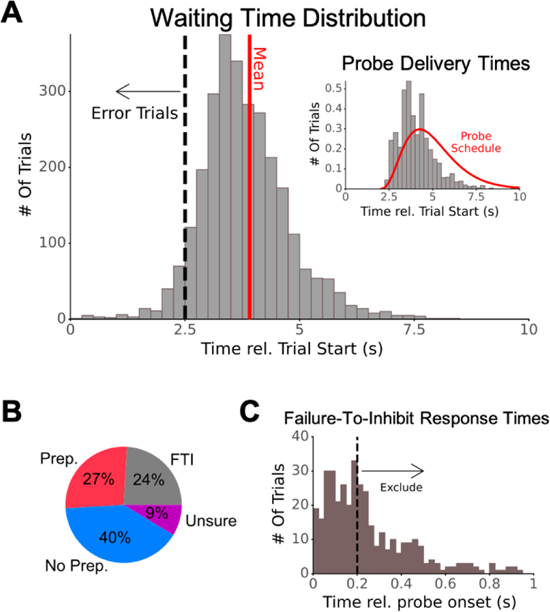
Behavioral results. **A**: Histogram of waiting times pooled across participants (for visualization. purposes only). Dotted line: 2.5 second cut-off for excluding responses from the analysis. Solid line: mean of waiting times (after excluding waiting times < 2.5 s). Inset: Histogram of probe delivery times (in gray; density; also pooled across participants) and the gamma distribution used for random probe delivery schedule (in red). **B:** Frequency of experimental conditions following the probe. Participants could answer the question about preparing to move with “Yes” coded as Preparation (Prep), “No” coded as No Preparation (No Prep), and “Unsure”. Or, if they moved within 200 ms of probe onset, the trial was categorized as failure-to-inhibit (FTI; see Methods for more information). **C:** Histogram of button-press times when participants moved following the probe, pooled across all participants. About 65.4% of movements were within 250 milliseconds of the probe, suggesting actions were already ballistic and participants were not able to inhibit their movements (a uniform distribution across 1 s after probe onset would predict 25% rather than 65.4%). However, we retained only those button presses within 200 milliseconds of probe onset, as that this was found to be the point of no return^37^.

In addition, for our ERP analysis, we were specifically interested in the RP. Therefore, to account for differences in underlying brain morphologies, we inspected the obtained ERPs aligned to movement onset for movement trials at electrodes Cz, CPz, FCz, and Fz. For each participant, we used data from the electrode where the EEG signal was most negative at the time of movement (using Cz for every participant reduced RP amplitude but did not meaningfully change our results). We omitted 4 participants (out of 24, 16.7%) who did not exhibit a visible RP at any of these locations. Therefore, our main inclusion criteria had 20 participants (Cz n=9, FCz n=9, Fz n=2), which is comparable to similar studies.

### 2.6 Statistical Analyses

The number of trials was not the same across participants (see Paradigm). Due to stochastic probe delivery and the dependency of reports on participants, our methods frequently result in different numbers of trials for different conditions (we split the probe-trial data into 4 cases; preparation reported, unsure, no preparation reported, and failure-to-inhibit). So, rather than traditional cluster-based ERP analysis, we opted to analyze single-trial data using mixed-effects models (LMEs, also called linear mixed-effects models). LMEs were implemented using the Pymer4 package in Python, which is a port of the commonly used Lme4 package in R. This correction was not necessary for time-domain EEG. In some cases, we also wanted to assess how data affected a binary variable (e.g. response to a probe being a ‘yes’ or a ‘no’). For these cases, we used a *logistic* mixed-effects model to assess the degree to which a given variable affects the log-odds of a binary outcome variable.

#### 2.6.1 Behavioral Analysis

We also used logistic mixed-effects models to assess whether participants were more likely to report that they were preparing to move if more time had elapsed since trial onset. To do so, we used the following regression equation with random slopes:

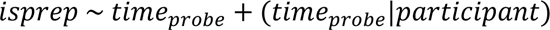

Where in the above, *isprep* is a binary response variable (1 for prep reported, 0 for prep not reported), and we regress time of the probe since trial onset (time_probe_) with a random intercept and effect of time for each participant. We complemented this analysis by regressing condition (Prep vs No Prep) on probe timing:

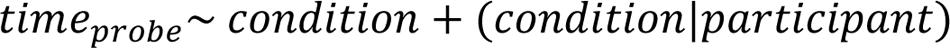

#### 2.6.2 Event-Related Potential Analysis

In Figure 3 we assessed the means of each ERP using linear mixed-effects models. For figure 3A, we assessed grand-average amplitude at each time point using the below equation with random intercepts (to avoid singularities in fitting, we omitted random slopes):

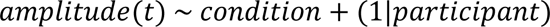

**Figure 3:**
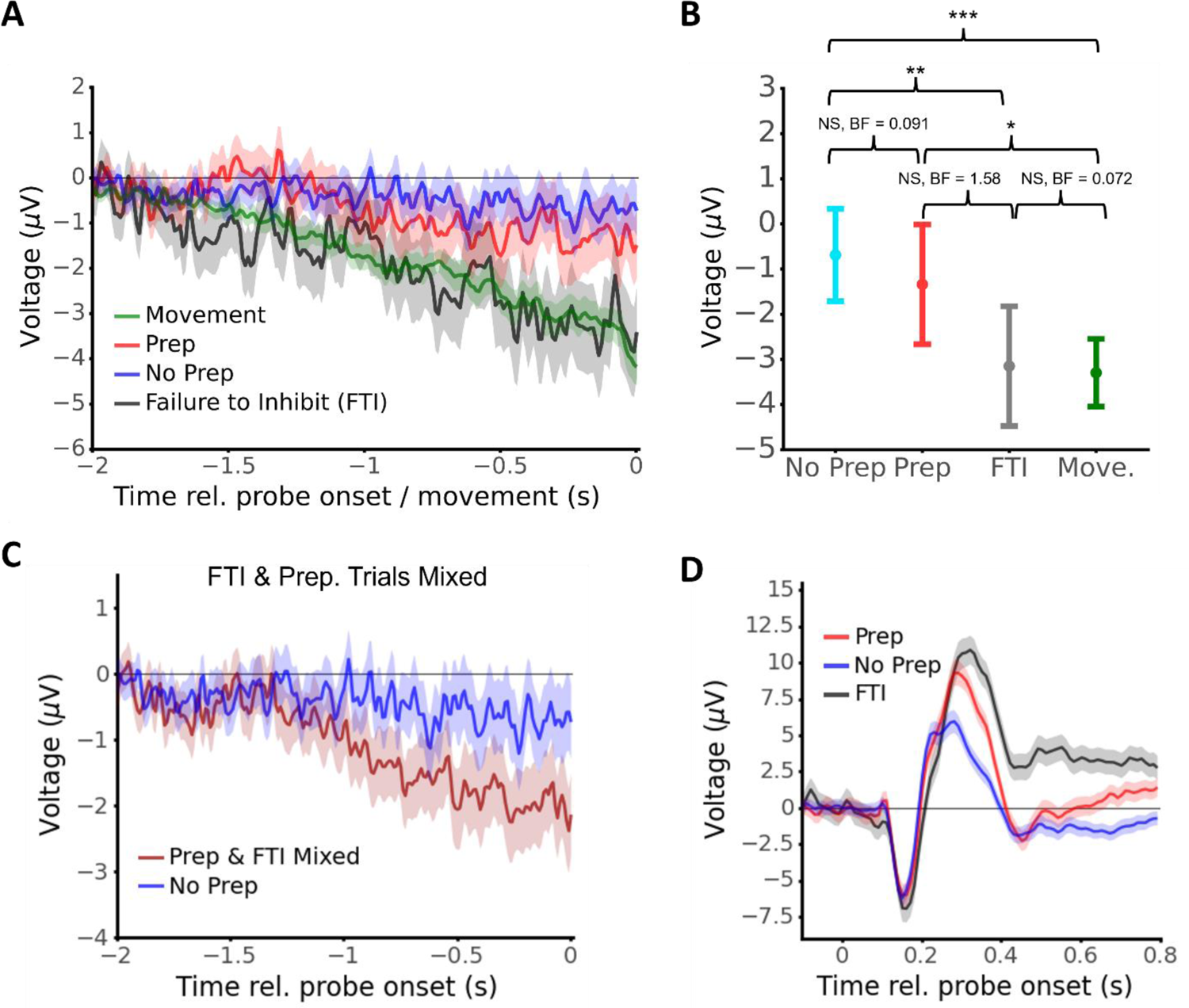
ERP Results: pre-probe RP amplitude is not associated with subjective reports of motor preparation. **A:** ERPs for Movement (green), Prep (red), No prep (blue), and FTI (black) trials. The ERPs are aligned to button-press onset for the Movement trials and to probe onset for other conditions. Solid lines are means and shaded regions are standard error (obtained via Linear Mixed Effects; LME). **B:** RP amplitude (mean & 95% confidence interval; LME analysis) in the 250 milliseconds before probe onset (Prep, No Prep, FTI) or movement onset (Movement). Statistical results from LME analysis post-hoc comparison (FDR corrected) & Bayesian ANOVA shown. **C:** Zoom on pre-probe/movement period for probe trials (means & SE; LME analysis). Separating FTI and Prep trials results in negative deflection only on FTI trials (left). But mixing FTI and Prep trials creates a downward, RP-like, deflection on “prep” trials (right). **D:** Post-probe ERPs aligned to probe onset (means & SE; LME analysis; baselined using period [−0.2,0.0] s relative to probe onset). All trials showed a negative deflection of similar magnitude followed by a positive deflection that reflected both whether participants failed to inhibit their movement as well as their later report.

**Figure 4:**
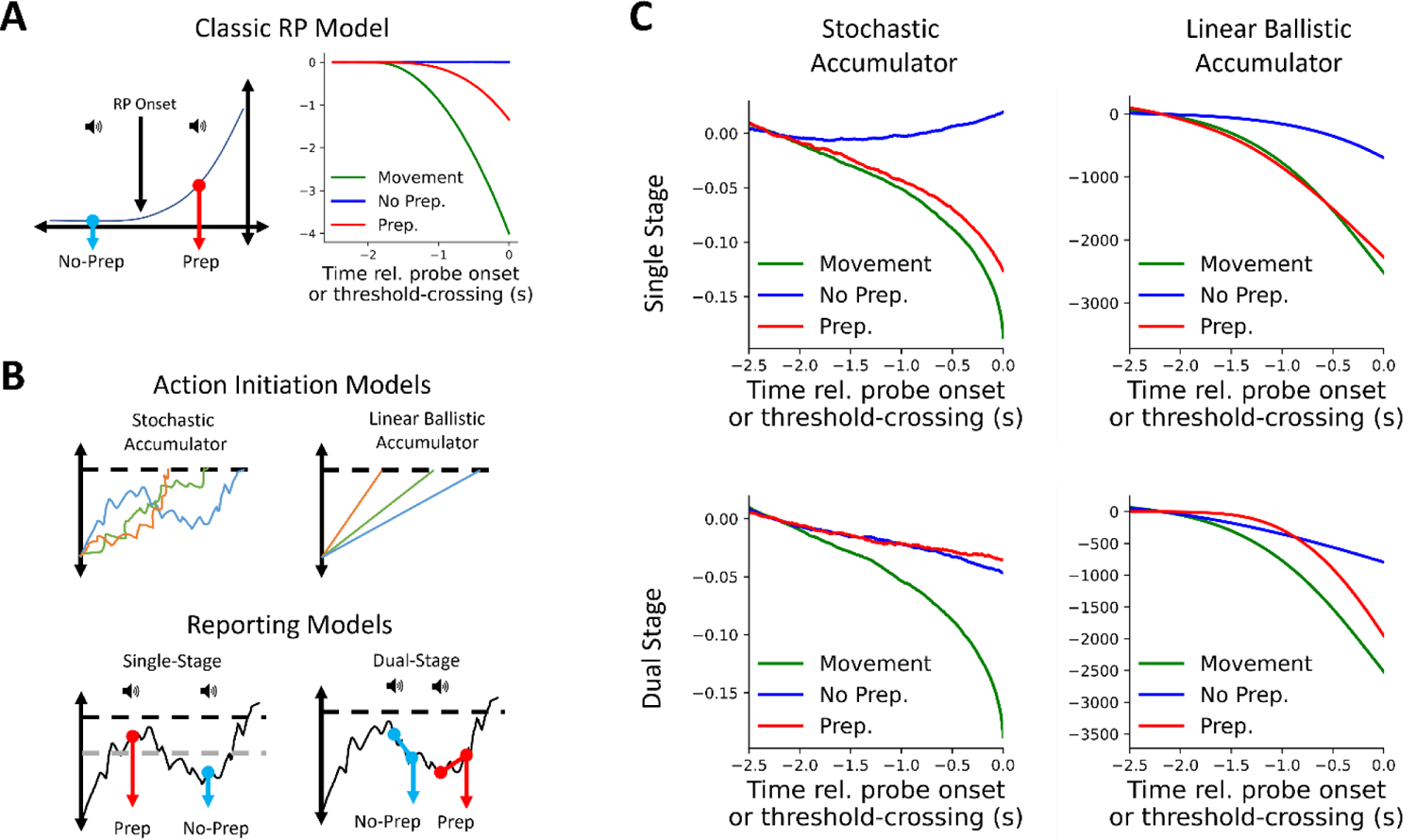
Computational modeling results: dual-stage metacognitive access to stochastic accumulation can explain ERP results. **A:** Classic model simulations. Left: model schematic. If the probe occurs before (after) an RP onset, the trial is categorized as a No Prep (Prep) trial. Right: modeling results. Prep trials show moderate deflection while No Prep trials do not. Move trials show stronger deflection. **B:** Modeling framework. Top: Schematic of two accumulator models; Left: stochastic accumulator: a noisy diffusion process triggers movement upon crossing a threshold (dashed line). Right: linear ballistic accumulator: linearly accumulating process triggers movement upon crossing the threshold. Bottom: Schematic of metacognitive reporting demonstrated on stochastic accumulator dynamics. Left: single-stage model, wherein reports are based on the magnitude of the accumulator process at the time of the probe (roughly, above or below the dashed gray threshold; see Methods). Right: dual-stage model, where participants’ reports are based on the *slope* of the accumulator immediately after probing. **C:** Modeling results. We tested two types of underlying accumulation dynamics each with two types of reporting models. With single-stage metacognition, both stochastic and linear ballistic accumulation predict differences between Prep and No Prep trials. Only dual-stage metacognition with stochastic accumulator dynamics matches ERP results, and also recreates the slight negative deflection before both Prep and No Prep trials.

The estimated average values obtained from the fitted LME models and their standard errors were used for Figure 3A. Figure 3C was obtained using the same model except re-coding FTI trials as Prep trials. Figure 3D was obtained using the same equation except with the EEG data baselined at (−0.2,0) instead of (−2.5,-2).

For assessing differences in pre-probe amplitude between No Prep, Prep, and FTI trials, we averaged single-trial EEG amplitudes in the 250 milliseconds before movement and used the following equation with random intercepts to assess differences between conditions.

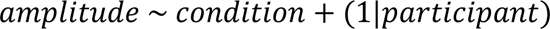

For the analysis of slope mentioned in-text, we used a similar equation except we did not average amplitude, instead fitting on the preprocessed EEG. We also included a random effect of condition to account for the relatively lower number of Yes trials present in our data (model would not converge with a random effect of time or the interaction term):

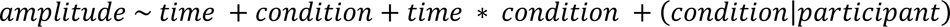

#### 2.6.3 Time-frequency analysis

For the analysis of power in the time-frequency domain in Figures 5A and 6, we fitted single-trial power data at every time-frequency pair using LMEs with a fixed effect of condition and random effect of participant (we did not use random slopes because of model fitting issues):

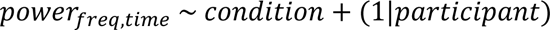

**Figure 5:**
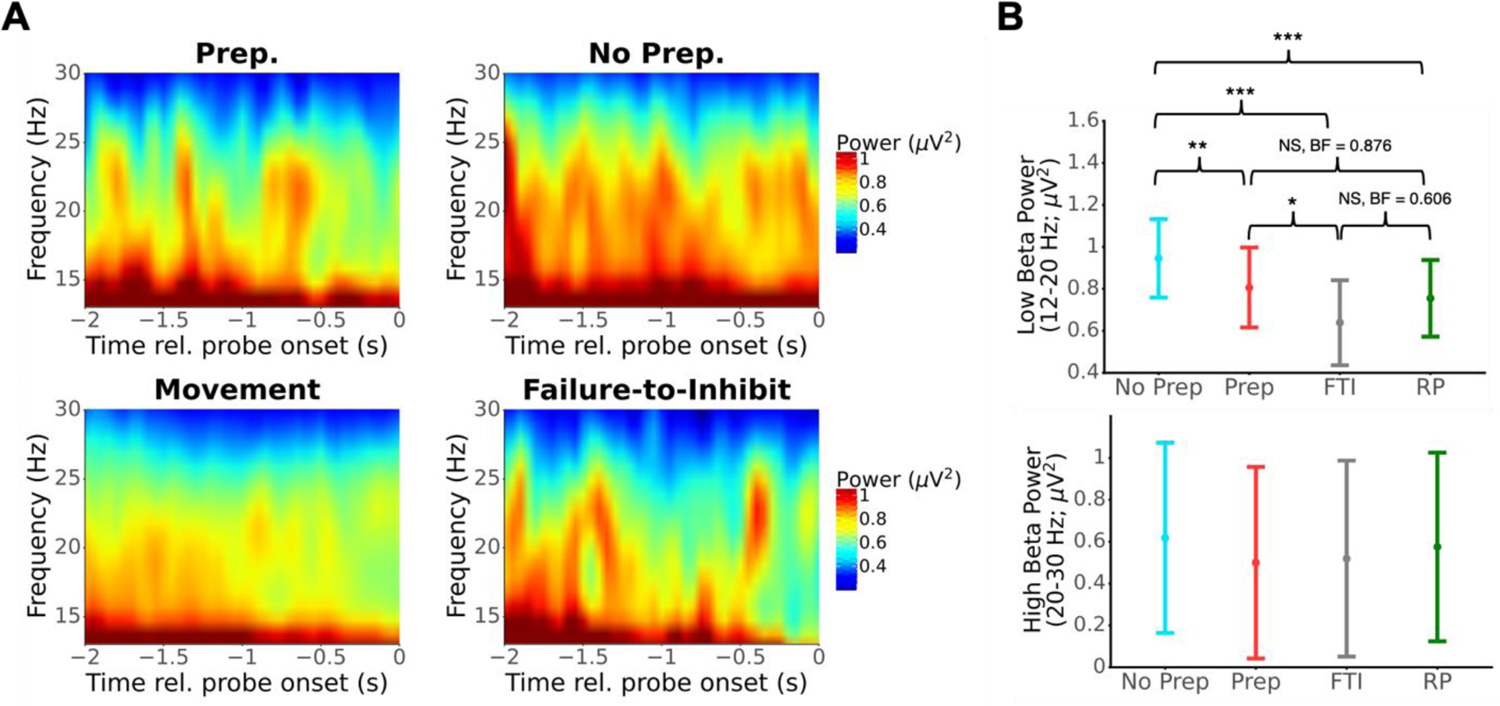
Analysis of pre-probe beta desynchronization. **A.** Spectrograms of beta-range activity (12-30 Hz) in the 2 seconds prior to probe onset (Prep, No Prep, Failure-to-Inhibit) or movement. Grand-average power estimates obtained by fitting an LME model at each time X frequency pair. **B.** Estimated beta power (mean & 95% CI, LME) in the 250 milliseconds before probe onset or movement for the low beta (12-20 Hz) and high beta (20-30 Hz) bands. No significant modulation was found in the high beta band, but the condition was significantly associated with pre-probe power in the low beta range. In particular, Prep trials were associated with lower pre-probe beta power than No Prep trials.

**Figure 6:**
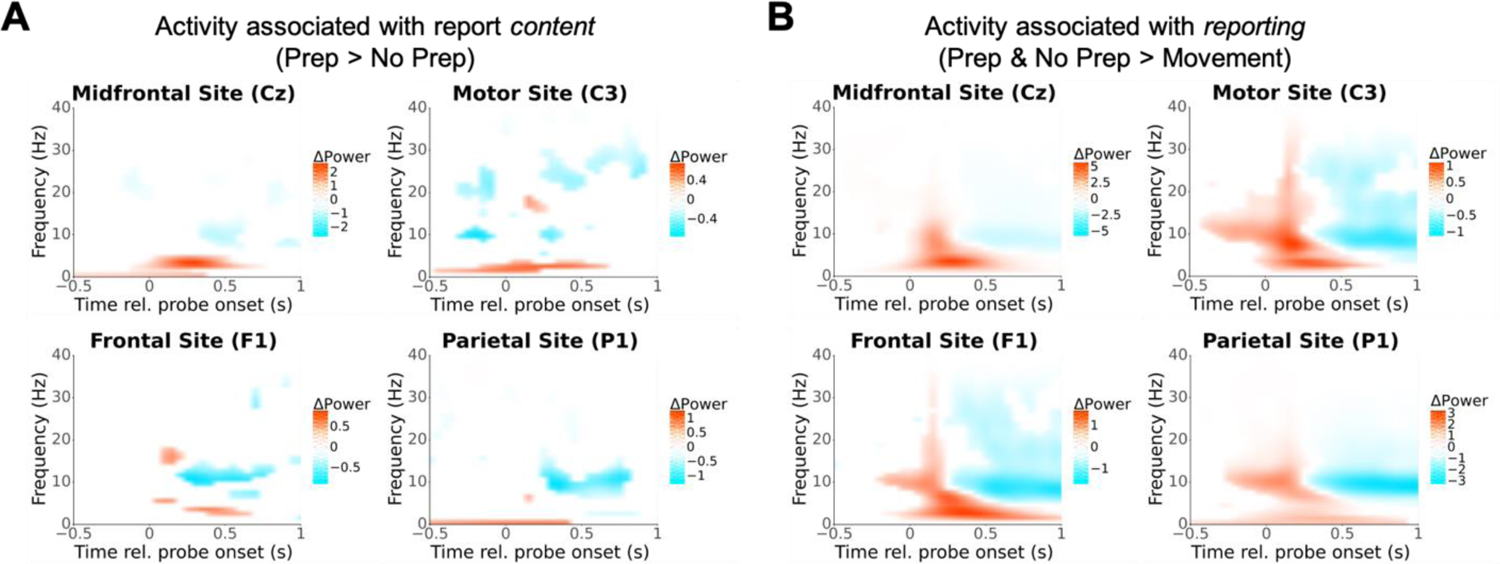
Results from exploratory time-frequency analysis. **A.** Time-frequency plots comparing report *content* at midfrontal, motor, frontal, and parietal sites (estimated difference in power between Prep vs No Prep trials estimated at each time X frequency pair using LME models). Red (blue) signifies greater power on Prep (No Prep) trials. Time-frequency pairs resulting in non-significant modulation are whited out (p > 0.05 uncorrected, less conservative because this is an exploratory analysis). Notably, report content significantly varies with activity in the second following probe onset. **B.** Time-frequency activity associated with making probe reports compared to not (estimated difference in power on Prep & No Prep trials compared to movement trials). Analysis conducted as in A. Reporting is associated with broad changes in activity in theta & alpha bands following probe onset.

Then, reported power in Figure 5A was taken as the marginal estimate of power for each condition. For Figure 6, we restricted the conditions to those specified and then conducted the same analysis and reported estimated differences using post-hoc testing. In Figure 6 we omitted time-frequency pairs that had an uncorrected p-value greater than 0.05 because that analysis was exploratory. Because these analyses entailed 1200 tests (one for each time-frequency pair) for each of four channels of interest, there are likely some spurious findings.

For specific analysis of pre-movement and pre-probe beta power (Fig. 5B) we averaged beta power across the low (12-20 Hz) and high (20-30 Hz) beta bands in the 250 milliseconds before probe/movement to get single-trial beta power values. Then, we fit an LME to these data using the following equation, including condition as a fixed effect and a random effect of participant:

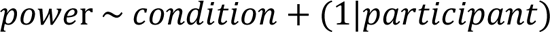

We established significance using post-hoc tests implemented in the pymer4 python library. For Bayesian analysis we ran a Bayesian ANOVA with the same structure in JASP^38^ version 0.6.13 and reported Bayes factors obtained from post-hoc tests.

### 2.7 Computational modeling

We also conducted computational modeling of the action-generation and metacognitive process to explain the basis of our ERP results. Note that all units are arbitrary and inverted for plotting to match the ERPs. In some cases, model parameters are taken from prior studies (e.g. stochastic accumulation). In those cases, we note the source of parameters. In other cases, model parameters were tuned manually to simultaneously obtain an RP shape and response time distribution close to real data on movement trials (i.e. without investigating differences based on Prep or No Prep trials).

#### 2.7.1 Classic RP Model

The so-called ‘classic’ interpretation of the RP has not been given a formal model before, so we began by doing so. The interpretation posits that RP onset corresponds to an ‘unconscious decision’ followed by a monotonic rise in activity. We implemented this in an accumulator framework as a noisy fluctuation process that does not cross threshold itself. After some time has passed (modeled as a gamma distribution), a strong input turns on (reflecting an unconscious decision) that drives the accumulator to the threshold in roughly 1 second.

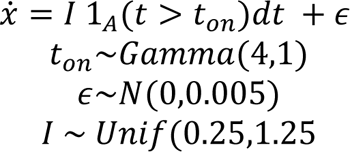

Where 1_A_(*u*) is the indicator function for *u* (i.e., gives 1 if *u* is true, 0 if *u* is false), and t_on_ is a randomly generated time at which a strong input drives the accumulator to the threshold. In our simulations, the threshold is 4/3 and dt=0.001. Individual runs, threshold-crossing distribution, and threshold-locked averages are shown in Fig. S2.

Probe delivery times were modeled as a shifted gamma function:

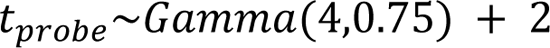

We first simulated 10,000 runs of 20 seconds, and then for each run calculated when t_probe_ occurred relative to t_on_. If t_probe_ occurred before t_on_, then the trial was categorized as a No Prep. Otherwise, if t_probe_ occurred after t_on_ but before threshold crossing, the trial was categorized as a Prep. If t_probe_ occurred within 200 milliseconds of threshold crossing, the trial was categorized as FTI. Otherwise, the trial was categorized as a Movement trial. Figure 5A bottom gives the average traces of these four conditions.

#### 2.7.2 Stochastic Accumulator Model

In the stochastic accumulator model, movement is triggered upon autocorrelated fluctuations (implemented as drift-diffusion process) crossing a threshold. We implemented the model following Schurger et al., 2012, using the equation:

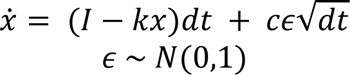

Where I is the drift rate, k is the leak, x is the accumulator amplitude, and c is a noise scaling term. We used the parameters (I = 0.11, k = 0.5, c=0.1, dt = 0.001, and a threshold of 0.298, from Schurger and colleagues^9^). Individual runs, threshold-crossing distribution, and threshold-locked averages are shown in Fig. S2.

#### 2.7.3 Linear Ballistic Accumulator Model

Recently, Bogler and colleagues^17^ suggested that linear ballistic accumulation dynamics^18^ could explain the early rise and shape of the RP. In linear ballistic accumulation, movement is triggered when a linearly increasing accumulator crosses a threshold. We therefore also simulated these dynamics to test whether our study’s findings could distinguish between the two underlying processes. In order to more accurately recreate the shape of the RP, we included a fixed delay of 2500 ms, such that the LBAc would only begin 2500 milliseconds into the trial. Note that without this nonlinearity, the average trace of the LBA before a threshold crossing would be a straight line. Also note that we tested a variety of parameter combinations, and they largely did not change our results. Our simulation was governed by the following equation (adapted from^17^):

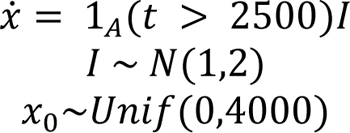

Where I is the slope (normally distributed), and x_0_ is the starting point of the accumulator. We used a normal distribution with mean = 2, standard deviation = 1 for slopes in our model whereas Bogler and colleagues^17^ used a normal distribution with mean = 1, standard deviation = 2. We made this change because, in simulating the model runs, we found that using their distribution would sometimes result in very shallow or even negative slopes on some trials. This led to long waiting times and positive-trending pre-probe EEG on No Prep trials, so we modified parameters to better match behavioral and EEG data.

#### 2.7.4 Pink-Noise Accumulator Model

Schurger^10^ hypothesized and found evidence that the RP might reflect a noisy autocorrelated *input* to an accumulator. This claim was evidenced by that RP amplitude being higher for trials that had shorter waiting times in comparison to longer waiting times. However, we did not find such a difference (Fig. S1B). Nevertheless, we included this model here as an alternative to the stochastic accumulator model. Briefly, we generated traces of pink noise with a 1/f slope of 1.5 using the “colorednoise” package in Python. If we let such traces be denoted P, our accumulators obeyed the following equation:

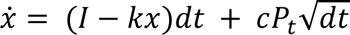

Where P_t_ denotes the t^th^ entry in P. We used the same parameters as the stochastic accumulator above, except we used the noisy input P as the traces for further analyses.

#### 2.7.5 Single-Stage Reporting Model

In our single stage reporting model, whether a trial was designated Prep or No Prep was based on accumulator amplitude at the time of the probe. Probe delivery times were generated using the following distribution:

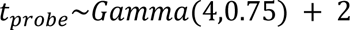

We first simulated 10,000 runs of 20 seconds, and then for each run calculated when t_probe_ occurred relative to threshold-crossing. Trials where t_probe_ occurred before threshold crossing were counted as probe trials. Otherwise, the trial was counted as a regular press trial.

The amplitude at the time of the probe was then extracted for each probe trial and categorized mirroring the percentages present in our behavioral data; the lowest 53.2% were categorized as No Prep trials, the next 11.4% were counted as Unsure and discarded, and finally the highest remaining 35.4% were counted as Prep trials. Using a median-split to define Prep versus No Prep, or using an 40-20-40 split for Prep, Unsure, and No Prep left results unchanged. Model runs were then grouped according to this classification and averaged for Figure 4C top row.

#### 2.7.6 Dual-Stage Reporting Model

In our dual-stage reporting model, a trial was designated Prep or No Prep according to the accumulator slope immediately following the probe, as opposed to the amplitude at the time of probe delivery. This model was developed following post-decision metacognitive models in perceptual decision-making^39–41^. In such models, confidence reports are based on whether evidence continues to accumulate in the same direction after reaching a decision threshold—if the accumulation continues, the participant is confident, but if the evidence stops accumulating the participant is less confident (some variants include whether the post-decision accumulation hits a secondary, higher threshold). We used a simplified version of such models by taking the slope of the post-probe accumulation. Probe delivery times were generated using the same equation as the single-stage model, and the simulations followed the same protocol. For trials categorized as probe trials, we fit a linear regression on the 200 milliseconds following the probe. We tested a variety of window sizes—100, 200, 300, and 400 ms—and the choice of window size did not substantively affect our results. So, we report results from the 200 ms window below, a duration that is in line with the perceptual-metacognition literature. Then, we categorized trials using the same percentiles as the single-stage model (i.e. highest post-probe slope percentile gets counted as Prep, lowest as No Prep, and medium as Unsure). Model runs were then grouped according to this classification and backaveraged for Figure 4C bottom row.

## 3 Results

Participants carried out a self-paced movement task, in which they were sometimes interrupted by an auditory tone that was scheduled to be delivered at a random time in each trial (the probe; see Fig. 1 and the Paradigm subsection of the Methods).

Participants were instructed to move whenever they felt like it after around 3 seconds had elapsed from trial onset, but to abstain from moving if the probe occurred. If they moved before 2.5 seconds had elapsed, they were given a warning, and the trial was aborted and designated an *Error trial*. Alternatively, if they moved after 2.5 seconds elapsed but before the probe went off, the trial was designated a *Movement trial* and participants immediately advanced to the next trial. Otherwise, the probe occurred before participants spontaneously moved. Despite being instructed not to move if the probe went off, participants sometimes pressed the button after probe onset. These presumably reflected failures to inhibit the movement (FTI) for those trials—i.e., trials where the participants could no longer inhibit the movement at probe onset (or, in other words, the movements were already ballistic). We designated trials where the button presses took place within 200 ms of probe onset as FTI trials because 200 milliseconds before movement marks the ‘point of no return’ at which action initiation processes become ballistic^37^ (see the Data Retention section of the Methods). Finally, if no movement followed the probe, participants were asked whether they were preparing to move when the probe went off. They could respond Yes, in which case the trial was designated a *Preparation trial*; Unsure, in which case the trial was discarded; or No, in which case the trial was designated a *No preparation trial*. The primary trials of interest were (1) Movement Trials (Move), (2) FTI trials, (3) Preparation (Prep) trials, and (4) No Preparation (No Prep) trials. Other trials were not further analyzed.

### 3.1 Behavioral results

On average (grand averaged across participant-specific averages), participants waited 3.98 seconds before moving (STD: 0.96, range: 3.38-5.91; see Fig. 2A for all waiting times pooled across participants) and were interrupted by probes on 39.2% of trials (STD: 18.2%, range: 14.0%-85.0%; see Fig. 2A insert for all probe delivery times). The heterogeneity across participants in the frequency of being probed is likely because some participants tended to move earlier (later) in the trial than others. Such movements were less (more) likely to encounter a probe before they moved. This also explains why the empirical distribution of probe delivery times was skewed earlier than the distribution used for probe scheduling (red line in Fig. 2A insert): later probes were more likely to be preempted by a participant’s movement and hence less likely to go off.

Among probe trials, participants reported preparation in 26.9% of trials, no preparation in 40.4%, and being unsure in 8.7%; a further 24.1% of trials were FTI (Fig. 2B). Thus, in 88.5% of non-FTI trials participants were confident in their ability to report having prepared or not prepared to move at probe onset, suggesting that they were largely capable of performing the task as instructed. Overall, averaged across participants, we obtained 117.2 ± 35.4 (mean ± STD) Movement trials, 20.2 ± 17.2 Prep trials, 33.3 ± 25.4 No prep trials, 7.0 ± 7.0 trials where participants were unsure about preparing to move, and 15.7 ± 11.6 FTI trials. Notably, most button presses following the probe occurred within 250 milliseconds of probe onset (65% compared to 25% at chance level assuming a uniform distribution across 1 second; Fig. 2C), suggesting that participants strove to follow instructions and inhibit movements after probe onset.

Removing FTI and unsure responses from probe trials, we found that the probability of reporting preparation significantly increased with time from trial onset (β_time_ = 0.322, p = 0.001; Logistic Mixed Effects). Prep trials also tended to have longer wait times than No prep trials (p = 0.023; Linear Mixed Effects), providing further evidence that participants were able to accomplish the task successfully. Similarly, among all probe trials, the probability a participant would fail to inhibit a movement also increased significantly with time from trial onset (β_time_ = 0.447, p < 0.001; Logistic Mixed effects).

### 3.2 EEG and modeling results

#### 3.2.1 The RP does not reflect awareness of motor intention

We investigated pre-probe EEG during Movement trials, FTI, Prep and No prep trials. We used linear mixed-effects (LME) analysis to estimate voltage at each time point for each condition (Fig. 3A; see Fig. S1C-D for other estimation methods). A clear RP was visible for Movement trials (Fig. 3A, in green; see individual participants in Fig. S1A).

FTI trials were also preceded by a significant negative deflection compared to Prep and No Prep trials, but we did not observe a difference between pre-probe EEG when comparing Prep and No Prep trials. To investigate the significance of these results, we investigated single-trial amplitudes in the 250 milliseconds before probe onset (or button-press for movement trials) using linear mixed-effects models (LME; implemented in python using the pymer4 library version 0.7.7^42^) and a Bayesian ANOVA (implemented in JASP^38^ version 0.6.13). Critically, we found no difference between pre-probe EEG amplitude for Prep and No Prep trials, with Bayes factors providing strong evidence that the two trial types were equivalent (Fig. 3B; post-hoc tests from LME analysis: No Prep vs Prep: t(3305.9) = 0.874, p_tukey_ = 0.004, BF_10_ = 0.091; No Prep vs Movement: t(2345.7) = 5.140, p_tukey_ < 0.001, BF_10_ =815.4; No Prep vs FTI: t(3206.6) = 3.260, p_tukey_ =0.006, BF_10_ = 21.0; Prep vs Movement: t(2903.4) = 2.948, p_tukey_ = 0.017, BF_10_ = 2.127; Prep vs FTI: t(2.994) = 2.059, p_tukey_ = 0.167, BF_10_ = 1.584; Movement vs FTI: t(3440.4) = −0.223, p_tukey_ = 0.996, BF_10_ = 0.072).

To account for concerns that the choice of baseline can artificially introduce differences in amplitude, we also investigated the EEG slope in the second before movement, because slopes are not affected by the choice of baseline. To do so, we fit LME models to the EEG data in the second before probe onset/movement. Consistent with the above analysis, pre-probe EEG slope was more negative in FTI and Movement trials compared to Prep and No Prep trials, whereas the slope for Prep and No Prep trials was not significantly different (post-hoc tests from LME analysis: No Prep vs Prep: Z= − 0.572, p_tukey_ = 0.940; No Prep vs Movement: Z = 6.657, p_tukey_ < 0.001 No Prep vs FTI: Z = 3.260, p_tukey_ < 0.001; Prep vs Movement: Z = 2.948, p_tukey_ < 0.001; Prep vs FTI: Z = 2.059, p_tukey_ < 0.001; Movement vs FTI: Z = 0.704, p_tukey_ = 0.896). However, pre-probe slope was indeed slightly negative on both Prep and No Prep trials (Prep: β_slope_ = −0.509 μV/s, 95% CI: [−0.935, −0.083]; No Prep: β_slope_ = −0.661 μV/s, 95% CI: [−0.962, −0.361]).

Notably, these findings were similar when we tested our data with more stringent exclusion criteria for participants (see Table S1).

Furthermore, we hypothesized that combining EEG signals from trials that included ballistic button presses (i.e., FTI trials) with EEG signals from trials that include presses stemming from actual metacognitive decisions that the participant was preparing to move (i.e., Prep trials) would result in an average ERP trace that would show a more negative deflection than No Prep trials. We tested this in our dataset and indeed found that combining Prep and FTI trials in this manner resulted in greater pre-probe negativity (Fig. 3C). Additionally, when investigating post-probe ERPs, we observed a negative deflection that began around 150 milliseconds after probe onset (Fig. 3D). This deflection was of similar magnitude across Prep, No Prep, and FTI trials. This initial deflection was followed by a positive deflection that was visibly largest and peaked latest for FTI trials, followed by Prep trials then No Prep trials.

#### 3.2.2 Reports based on stochastic accumulation can explain ERP results

Our ERP results suggest that there is no difference in pre-probe RP amplitude across trials between the condition where preparation was reported and the one where it was not, though our behavioral results suggest that participants were able to report on their state of preparation. For this to be possible, whichever feature of neural activity is accessed for these reports must be *statistically independent* (or only weakly dependent) from pre-probe RP buildup. In this section, we use computational models of the putative action-initiation neural mechanisms to investigate what processes could underlie our ERP results.

First, we looked at the classic RP interpretation—that its deflection reflects motor preparation following an unconscious decision to move^8^. Under that interpretation, probing after (before) the onset of the RP will result in the reported presence (absence) of motor preparation^24,26^. We developed a simple computational model to capture the essential components of this interpretation. Briefly, activity fluctuates noisily (random walk) until the onset of an “implicit decision”, after which a strong input drives activity up towards a threshold. Movement is initiated when that threshold is crossed. Although the RP usually has an exponential shape, we used a linear input for simplicity (see Classic RP model in the Computational Modeling section of the Methods; results are the same if we use an exponential buildup). In this model, if a randomly timed probe is delivered before (after) the implicit decision, the trial would be categorized as No Prep (Prep) (Fig. 4A left). If the probe was delivered after threshold-crossing, the trial was designated a Movement trial (we did not model FTI trials for simplicity). In this framework, pre-probe activity is statistically dependent on report (by definition), because the report explicitly depends on whether not the “implicit decision” to move has been made. Accordingly, we predicted that pre-probe activity would differ across Prep and No Prep trials for the classic RP model.

We simulated 10,000 trials under these conditions (Fig. 4A; see Classic RP Model section of the Supplementary Methods for details; see Fig. S2 top for example runs). Consistent with our predictions, we observed a flat ERP on No Prep trials (blue), a slight negative pre-probe deflection in Prep trials (red), and a stronger negative deflection before the probe or movement on FTI and movement trials (black and green, respectively). Comparing the model’s predictions to our empirical results (Fig. 3A) suggests that they are qualitatively not a good fit. In particular, the model predicts that there would be a difference between Prep and No Prep trials that grows with time, which we did not observe in the empirical data. Hence, the classic RP model is incompatible with our empirical results.

We next considered two prominent computational models of action initiation--stochastic accumulation^9,11^ (Fig. 4B top left) and linear ballistic accumulation^17,18^ (LBAc; Fig. 4B top right). In stochastic accumulation, movement is initiated when a noisy diffusion process crosses a threshold (see Computational Modeling in Methods for details).

Critically, that process is not monotonic—it can rise and fall until threshold-crossing—and only resembles an exponential rise when each run is aligned to its threshold crossing and many trials are then back averaged^9^. In our linear ballistic accumulator model, movement is initiated when a monotonic linear signal that begins increasing at trial onset crosses a threshold. The accumulator’s slope is drawn from a half-normal distribution at the onset of the accumulation process, and the accumulation then proceeds linearly & monotonically with that slope (see Fig. S2 for individual model runs).

We combined these two action-initiation models with two models of the reporting process. In the first reporting model reports were based on the accumulator amplitude at the time of the probe: if the accumulator is relatively high (low) at probe onset—i.e., if the accumulator was closer to (farther from) the threshold—the trial was categorized as Prep (No Prep) (Fig. 4B bottom left, where—schematically—the gray dashed line marks the threshold between preparing and not preparing; see Single-Stage Reporting Model in Methods for details). We termed this a single-stage reporting model. Notably, both stochastic and linear ballistic accumulation are autocorrelated processes, and thus amplitude at a particular time (e.g. at probe onset) is statistically dependent on the activity immediately prior. Therefore, we predicted differences in pre-probe RP for both models when accumulator amplitude served as the basis for reports. Hence, we also considered a second reporting model based on the *direction* of the post-probe accumulation. If the slope of the accumulator was relatively high (low) in a short window following probe onset, the trial was categorized as Prep (No Prep) (Fig. 4B lower right; we used 200 milliseconds for presented simulations but tested a variety of windows and window size did not qualitatively change our results). We termed this a dual-stage reporting model (see Single-Stage Reporting Model and Dual-Stage Reporting Model in Methods for details). Perceptual metacognition is sometimes better explained by dual-stage models, where reports are based on whether evidence continues to accumulate after crossing the decision threshold^39,40^, suggesting such a metacognitive model may be appropriate in the domain of self-initiated actions. Critically, because stochastic accumulation is a summation over white noise, whether the accumulator increases or decreases at any time point is only weakly independent from prior activity (with said weak dependence emerging due to accumulator input and leak). Therefore, we predicted no difference between pre-probe activity for Prep and No Prep trials when combining stochastic accumulation with dual-stage reporting. During linear ballistic accumulation, however, accumulator trajectory is linear, so slope at any time is not independent from previous activity, and thus we predicted differences in pre-probe RP for this model.

We therefore simulated two accumulation models, each with two reporting models, hence four models overall: (1) stochastic accumulation with single-stage reporting, (2) stochastic accumulation with dual-stage reporting, (3) linear ballistic accumulation with single-stage reporting, and (4) linear ballistic accumulation with dual-stage reporting. We simulated 10,000 runs of each model type and contrasted average traces before threshold-crossings and simulated probe onset. Both stochastic accumulation and linear ballistic accumulation recreated the shape of the RP in Movement trials (inverted to match the negative trajectory of EEG; Fig. 4C). Consistent with our predictions, model (2), stochastic accumulation with dual-stage reporting (bottom left in Fig. 4C), was the only model that was able to recreate our finding that pre-probe EEG traces on Prep and No Prep trials were largely overlapping (Fig. 3 top).This model also predicted a slight negative slope in pre-probe EEG on both Prep and No Prep trials, which was also observed in experimental data (Fig. 3). All other model combinations led to large visible differences in pre-probe activity when comparing Prep and No Prep trials (Fig. 4C).

Another version of the stochastic accumulator model, which we term the pink-noise accumulator, has been suggested to underly self-initiated action and the RP^10^. In this model, the RP reflects an autocorrelated noisy *input* to stochastic accumulation, rather than the state of the accumulator itself^10^. Interestingly, this variant of the stochastic accumulator predicted differences between Prep and No Prep trials for both single and dual-stage models (Fig. S3A). We also tested models where the metacognitive decision was made after a delay (to account for auditory processing and task-shifting, among other possibilities), and the results were largely the same as the models discussed here (Fig. S3B).

Of the computational models we tested stochastic accumulation with dual-stage reporting was the closest to recreating our ERP results. In stochastic accumulation, This is because slope following probe onset (the feature used in dual-stage reporting) has a weak statistical relationship to pre-probe activity, whereas the features used for reporting in all other models were strongly related to pre-probe activity. Hence, this is the only model that shows little-to-no difference in pre-probe activity when comparing Prep and No Prep trials. However, another possibility is that reports of preparation are made based on a neural feature unrelated to the RP, as has recently been suggested^25^ in opposition to prior probe method studies^23,26,27^. We assessed this possibility by repeating the previous modeling and classifying probe trials as Prep or No Prep trials randomly (as would be the case if the neuronal basis of reports was unrelated to the RP). As expected, we found no difference in the pre-probe RP amplitude for all tested models under this assumption (Fig. S3C).

#### 3.2.3 Pre-probe beta power may track reported state of preparation

We next investigated changes of beta power in contralateral motor cortex (electrode C3) before movement & probe onset. We observed beta desynchronization on Movement and FTI trials (Fig. 5A). We also observed visibly less beta power prior to probe onset for Prep compared to No Prep trials. We next computed average beta power in the 250 milliseconds (as in our RP analysis) prior to probe onset / movement to investigate whether differences between conditions were significant. We separately analyzed low beta (12-20 Hz) and high beta (20-30 Hz) because pre-movement desynchronization was most prominent in the low beta band in our data (analyzing beta as a whole, 12-30 Hz, led to similar but weaker results as analyzing the low beta band). Furthermore, pre-movement desynchronization occurs at different times for low and high beta band activity^7^, suggesting separate analysis of these bands is appropriate. We fit LME models and conducted post-hoc tests to investigate differences and used a Bayesian ANOVA in JASP to estimate Bayes Factors. Pre-probe power in the low beta-band significantly differed across trial type (Fig. 5B upper; No Prep vs Prep: t(3979.6) = 3.421, p_tukey_ = 0.004, BF_10_ = 0.711; No Prep vs Movement: t(3987.4) = 6.459, p_tukey_ < 0.001, BF_10_ > 1000; No Prep vs FTI: t(3974.6) = 5.591, p_tukey_ < 0.001, BF_10_ > 1000; Prep vs Movement: t(3982.4) = 1.467, p_tukey_ = 0.458, BF_10_ = 0.876; Prep vs FTI: t(3975.9) = 2.857, p_tukey_ = 0.022, BF_10_ = 60.2; Movement vs FTI: t(3972.8) = 2.287, p_tukey_ = 0.101, BF_10_ = 0.606). However, we found no differences in pre-probe power in the high beta-band across conditions (Fig. 5B lower; No Prep vs Prep: t(3974.3) = 1.987, p_tukey_ = 0.193, BF_10_ = 0.066; No Prep vs Movement: t(3978.0) = 1.003, p_tukey_ = 0.748, BF_10_ = 0.046; No Prep vs FTI: t(3972.3) = 1.233, p_tukey_ = 0.606, BF_10_ = 0.097; Prep vs Movement: t(3975.5) = −1.442, p_tukey_ = 0.473, BF_10_ = 0.057; Prep vs FTI: t(3972.8) = − 0.222, p_tukey_ = 0.996, BF_10_ = 0.106; Movement vs FTI: t(3971.6) = 0.750, p_tukey_ = 0.877, BF_10_ = 0.180).

#### 3.2.4 Exploratory analysis of what neural activity relates to reporting and report content

We next investigated other potential relations between neural activity and the reported awareness of motor preparation. We focused on electrodes above brain regions that have previously been associated with intention awareness—frontal (F1), central (Cz), contralateral motor (C3—as participants always moved their right hand), and parietal (P1) sites. We extracted power at different frequencies and times using a Morelet wavelet decomposition^36^ (see Time-Frequency Decomposition in Methods). We assessed whether spectral power depended on trial type by fitting a LME model to each time-frequency pair. Figure 6 shows the results of this exploratory analysis, with time-frequency pairs that had p > 0.05 (uncorrected) whited out. These analyses were exploratory and, in the latter case, likely confounded by factors such as auditory processing, yet we thought they may be of interest to the field and opted to include them.

To investigate what activity reflected the *content* of reports, we compared Prep to No Prep trials. Activity in several sites seemed to associate with report content (Fig. 6A). Notably, much of the modulation we found was present after probe onset. Most prominently, we found midfrontal delta (0.5-4 Hz) and theta (4-7 Hz) power was higher (red) on Prep than on No Prep trials immediately after probe onset (Fig. 6A top left; peak modulation at 2.52 Hz, 350 ms after probe onset, p = 3.44 x 10^-28^ uncorrected). We also found a cluster of decreased (blue) parietal alpha (7-12 Hz) power (Fig. 6A top right; most extreme p = 4.04 x 10^-8^ at 9.6 Hz, 350 ms after probe onset). Notably, C3 was the only site we investigated where we observed differences prior to probe onset, likely reflecting the pre-probe beta differences we investigated in Figure 5.

To investigate what activity reflected the *metacognitive processes involved in reporting* we compared Prep and No Prep trials together to Movement trials. Once again, activity in several sites seemed to associate with the act of reporting (Fig. 6B), while modulations largely occurred after probe onset, excepting beta-band activity over contralateral motor cortex, which likely reflects the stronger beta desynchronization on movement trials compared to Prep and No Prep trials (hereafter just referred to as “Probe trials”). We found a cluster of broadly increased theta power following probe onset on Probe trials compared to Movement trials (Fig. 6B top left; peak modulation at 2.52 Hz, 350 ms after probe onset, p = 3.44 x 10^-28^ uncorrected), which may be an artifact of hearing the probe. We also found a cluster of decreased alpha power beginning several hundred milliseconds after probe onset, most prominently over Parietal sites (Fig. 6B bottom right; peak modulation at 13.67 Hz, 600 milliseconds after probe onset, p = 4.48 x 10^-23^ uncorrected). Again, these analyses are highly exploratory and it is not clear whether these results reflect the reporting process per se or another process, due to our design.

## 4 Discussion

We set out to investigate what process the readiness potential (RP) reflects and how it relates to the awareness of the preparation to move. We ran a variant of a self-initiated action task, where we sometimes interrupted participants before they moved with an auditory probe. We instructed participants to inhibit movement in response to the probe and only then report whether they were preparing to move when the probe arrived. This modification allowed us to separate cases of late-stage action initiation, when movement was already ballistic and participants failed to inhibit it, from earlier stages.

Contrary to prior results^23,24^, we found no relation between reported awareness of motor preparation and pre-probe RP-like buildups after controlling for trials where the participants failed to inhibit their movement. As we showed, such ballistic actions were preceded by an RP. This finding suggests that prior reports that pre-probe RP-like deflections were associated with reported awareness of motor preparation reflect the late but not early RP.

Our findings suggest that the early buildup of the RP does not reflect a phenomenal experience of motor preparation or of initiating an action. These results are therefore in line with findings that the timing of the RP’s buildup does not relate to reported timing of the conscious intention to move timed using a clock^5,21,22^. Our results are also in line with another recent probe study, which found similar results regarding the RP^25^. In the same vein, our findings also suggest that prior claims that the pre-probe RP amplitude relates to awareness of motor preparation are based on the inability of some probe paradigms to tease apart ballistic movements (i.e., in which the probe was delivered when the participants could no longer refrain from moving) from trials where the participants report preparing to move. More specifically, Parés-Pujolràs and colleagues^23^ found an RP before probes when participants reported they were preparing to move. However, in their paradigm, participants were instructed to move immediately if the probe occurred when they were already preparing to move (in contrast to our instruction to participants *not* to move in such cases). Hence, due to their experimental design, it was impossible for the authors to tease apart trials where the probe arrived when movement was already ballistic from trials where the participants moved in response to the probe to indicate they were preparing to move. Similarly, Schultze-Kraft and colleagues^24^ found that participants were more likely to report having an intention upon being probed when an online, real-time, closed-loop decoding system determined that an RP was present. However, predicting when someone will make a spontaneous movement from EEG is notoriously difficult: for instance, Bai and colleagues^43^ were able to predict an upcoming movement only around 0.62 seconds before its onset on average (with an average accuracy of 78% correct; other studies found similar results^44,45^). This is while RP onset occurs more than 1.5 seconds before the movement. Notably, those studies achieved their results by simultaneously analyzing tens of electrodes. Therefore, Schultze-Kraft and colleagues’ finding that probes delivered when a BCI detected an RP were more likely to elicit reported awareness of motor preparation were also likely based on the later stages of the RP.

Parés-Pujolràs and colleagues^25^ recently ran another probe method study that controlled for ballistic button presses by having participants report whether they were preparing to move at probe delivery using the hand opposite the one used for the self-paced action task. With this change in task design, they found no difference in pre-probe RP amplitude when comparing trials in which participants reported preparing versus not preparing to move. They hypothesized that their prior results^23^ were “confounded by some trials where self-paced action preparation was interrupted prior to awareness.” Our results suggest that it was not just that action preparation was interrupted, but rather that participants were at the very latest stages of action initiation when the probe was delivered and could thus no longer refrain from moving.

We next contrasted the ability of the classic RP model, stochastic accumulation, and linear ballistic accumulation to explain our RP results. Only stochastic accumulation, when combined with a dual-stage metacognition model, could successfully explain our EEG findings. Among other things, all other models predicted differences in pre-probe RP amplitude, contrary to our results. Critically, our ERP results directly contradict conceptual models wherein RP onset corresponds to a specific neurocognitive event, after which participants (if probed) will report that they were preparing to move^23,24,26,27^.

More generally, our computational-modeling results support the view that the RP reflects stochastic accumulation to a threshold that is aligned to and backward-averaged from threshold-crossings, and, furthermore, that subjective reports reflect dual-stage metacognitive access to this process in response to hearing the probe. Only this combination of action-initiation and reporting models successfully explained our ERP results. Furthermore, our model also predicted the slight negative deflection in pre-probe ERP, which we observed on probe trials regardless of later reported awareness.

Stochastic accumulation is a more recent yet influential model of self-initiated action and the RP^9–12^. Our results support stochastic accumulation over classic RP models^2,26^ or linear ballistic accumulation^17^. Surprisingly, all the models that were based on single-stage reporting—i.e., on accumulator activity at the time of the probe—failed to recreate our empirical results. Instead, our results suggest that dual-stage reporting underlies reports of awareness of motor preparation. Hence the findings are in line with work on perceptual metacognition, which suggests that post-decision evidence accumulation drives confidence reports^39,40^. Dual-stage reporting could conceivably be implemented by a separate population of neurons that has the capacity to monitor the accumulator, but only does so when triggered by the probe. Notably, stochastic accumulation with dual-stage metacognition (which was simulated using 10,000 trials—many more than is feasible to collect empirically) suggests that there may indeed be a slight difference between pre-probe neural activity when comparing Prep to No Prep trials (see Fig. 4C). However, that difference is too small to be resolved using EEG, given the limited number of trials that can be recorded empirically together with EEG’s low signal-to-noise ratio. Future intracranial work may be required to resolve this issue.

Notably, Schmidt and colleagues^14^ propose a general model of the RP (without defining it mathematically), wherein slow fluctuations in scalp EEG signal bias action timing. In their model, movements are more likely during the negative phase of infraslow cortical oscillations (<0.1 Hz). This model is fundamentally different from stochastic accumulator models, which assume that the RP reflects an accumulation process that begins at trial onset (although the two could possibly be merged, if the ongoing slow oscillations were fed into an accumulator—though this was beyond the scope of the current study).

Schmidt and colleagues hypothesized that infraslow oscillations would be related to the ebb and flow of a subjective feeling of “readiness” to move^14^. On one reading, our results provide evidence against that hypothesis: the lack of difference between pre-probe RP across Prep and No Prep trials suggests that the RP (whether it reflects slow cortical fluctuations or another phenomenon) does not reflect the subjective experience of *already* feeling readiness to move prior to being probed. However, on another reading, our results are compatible with Schmidt and colleagues’ hypothesis due to the specific phrasing we used in our study. We asked whether participants had already begun preparing to move, which is an active process that involves major shifts in neural state space^46^. However, Schmidt and colleagues specifically hypothesized that slow cortical fluctuations impact the *feeling of readiness* to move, which may be more of a passive process. Whether participants would treat these two questions the same or differently is an interesting open question for further study (although see^25^).

Our findings have major implications for the validity of the probe method for timing intention onset. The probe method was originally introduced as an alternative to the clock method for timing intention onset^26^. “T-Time”, as it was called by Matsuhashi and Hallett, was obtained by instructing participants to inhibit their movements in response to the probe *if they were already preparing to move*, and to ignore the probe otherwise. Hence, probes would result in vetoed movements when participants were preparing to move. So, the distribution of probe timings relative to movement onsets would have a “dip” in probe frequency at a certain time before movement corresponding to T-Time (see^26^ for details). Using this method, T-Time (usually 1-2 s before movement onset) was far earlier than estimates of intention onset from the clock method (usually around 200 ms before movement onset). This may even overturn the classic interpretation that the RP emerges before conscious intention^47^. However, the validity of this method relies on two crucial assumptions. For T-Time to be an accurate measure of intention onset, it requires that the assessment of whether one is preparing to move can be made (1) instantly, and (2) without affecting the movement generating process. If (2) is met but (1) is not, and assessing whether one is preparing to move takes, say, 300 milliseconds, then T-Times would be biased to be 300 milliseconds further from movement than the “true” timing of intention onset. If (2) is not met, the whole movement-generation process might be thrown off when participants report T-Time, making it less likely that T-Time captures the timing of a specific, meaningful event prior to movement onset, let alone specifically tracking the onset of intention. Furthermore, under the stochastic-accumulation interpretation of the RP at least, the process of accumulation over noise leaves no reason to expect events relevant to action initiation to reliably take place 1-2 seconds before movement onset. Indeed, under this interpretation the only events accompanying action initiation are trial onset, when the accumulator process is initiated, and threshold crossing, shortly before movement onset. There was an extension of the stochastic-accumulator model that incorporated a sub-threshold value, whose crossing would set of an intention to move^10^. Nevertheless, while probe studies offer valuable insight into the nature of conscious intention, our results and the stochastic accumulator model suggest that care should be taken when designing and interpretating such probe studies.

Our study is limited by our assumption that responses to the probe are made based on a feature of the RP (a common assumption in probe paradigms^23,24,26,27,48^). However, the critical requirement for our results is that the basis of reporting is weakly correlated with pre-probe RP buildup, which could also be satisfied if reports were based on a neural signal unrelated to the RP. Indeed, simulations showed that if trials were randomly classified as Prep or No Prep (as would be the case if they were based on a signal independent from the RP), it could lead to EEG results similar to those we observed under all the models we tested (Fig. S3C, though as discussed there, random guessing is unlikely due to low rates of participants reporting being unsure of their preparation, probability of reported preparation increasing with trial duration, and the other differences between Prep and No Prep trials we observed). Our results could thus be stated as the following: either (a) reported awareness of motor preparation is based on a feature of the RP, in which case only stochastic accumulation can explain our ERP results, or (b) reported awareness of motor preparation is not based on the RP but rather on another neural signal, such as beta power in contralateral motor cortex^25^. In either case, our study provides strong evidence against the influential conceptual model offered by Matsuhashi & Hallett^26^, wherein participants are initially “latently aware” of an intention to move, where the emergence of latent awareness is related to RP onset.

Recently, Parés-Pujolràs and colleagues^25^ suggested that beta oscillations, rather than the RP, relate to subjective reports of motor preparation. In line with their results, we found that pre-probe beta power over contralateral motor cortex was slightly higher for Prep trials compared to No Prep trials (Fig 5). Although these results suggest that beta power in motor regions, and not the RP, underlie awareness of motor preparation, it is not clear how motor-related beta power and the RP interact with each other. Notably, our results are similar to those from more traditional stop-signal paradigms^49,50^, in which participants are given a go-cue, which is sometimes followed by a stop-signal, signifying they should cancel their movement. From that perspective, our study could be considered a stop-signal paradigm without a specifically timed go-cue. In line with our results, stop-signal studies have found that beta-band activity is relevant for initiating and stopping actions^50–53^, suggesting that the pre-probe differences in beta power we observed may indeed be related to initiation processes. Taken together, our results therefore raise three critical questions for future studies on volition: (1) are reports in probe paradigms made using a feature of the RP or beta desynchronization? (2) can beta desynchronization prior to self-initiated movement be explained by a stochastic accumulator-type process? And, (3) how does beta desynchronization in the motor cortex relate to the RP? Although many studies have been conducted on the RP and beta desynchronization, few look at them simultaneously. Furthermore, although they have different cortical sources^7^, the motor cortex and SMA are strongly interconnected, and it is unlikely the two phenomena are completely independent.

We observed that post-probe oscillatory power in the theta and beta frequency bands was related to reported awareness of motor preparation. Interestingly, oscillations in those frequency bands have been observed during metacognition in other contexts^54–57^. But our paradigm differs from metacognition paradigms where participants reported confidence about a perceptual stimulus^39^ or the quality of a value-based decision^58^. It is therefore unclear whether these oscillations relate to metacognition per se. Theta oscillations have also been associated with cognitive control^59–61^, which may have been required to suppress movement when participants were already preparing to move— i.e., in Prep trials in our paradigm—leading to the increased theta power on Prep trials that we observed. Notably, it has been proposed that midfrontal theta oscillations broadcast a need for cognitive control^61^, and then inter-region communication via beta-band synchrony underlies metacognition^57^. Our results largely align with these suggestions; although our study was not properly controlled to make inferences about what specific mechanisms these power modulations reflect. Future research on metacognition and motor preparation could use a modified version of our paradigm—for example by including some trials where probes are delivered but participants are required to make a perceptual decision (such as whether the probe was of high or low pitch) rather than a metacognitive decision—to investigate that hypothesis more rigorously.

Our study was also limited in other ways. First, we required participants to inhibit their movement in response to the probe and only then report whether they were preparing to move. Hence, participants’ reports in our paradigm relied on working memory, which was part of the impetus behind probe studies^23,26^. This design therefore may introduce memory-associated noise and biases to participants’ reports, which were at least partially removed from fully online probe methods. However, importantly, even in such online probe studies, decisions about one’s underlying state cannot be made immediately: at the very least, one must perceive the probe and then decide whether one was already preparing to move, thus introducing some delays and relying at least to some degree on working memory. Furthermore, in the present study the timing of participants’ movements was arbitrary. Such actions are often habitual and could involve less overt awareness compared to actions made purposefully. Future research could increase the intentionality behind movements and increase participants’ attention during experiments, for example by providing reasons to move at one time point or another and rewarding adherence to task demands.

Taken together, our study parsimoniously resolves ongoing debates and conflicting findings regarding what kind of process underlies the RP and how that process relates to the conscious intention to move. Our ERP results suggest the RP is not directly related to the conscious experience of initiating action. But our computational modeling suggests that the neural process underlying the RP or beta desynchronization may nevertheless be accessible via metacognitive mechanisms. These results highlight the importance of investigating metacognitive mechanisms when studying conscious intentions. Consider that we are not constantly conscious of all our actions, decisions, and intentions when acting voluntarily—we can reach for a doorknob or pick up a cup of coffee without consciously representing, in advance or after the fact, all of the intentions that led to that action. Yet, we can introspectively access our intentions and report them if needed. We suggest that such metacognitive mechanisms are crucially involved in probe paradigms that target motor preparation and may be involved in accessing intention-related information. The version of the probe method that we developed thus offers an opportunity to study conscious intentions more generally, which is a long-standing goal of volition neuroscience.

## Data and Code Availability

Data and code will be made available upon publication of the article.

## Author Contributions

**Jake Gavenas:** conceptualization, methodology, software, formal analysis, investigation, visualization, writing – original draft; **Aaron Schurger:** methodology, resources, writing – review & editing, supervision; **Uri Maoz:** conceptualization, methodology, validation, resources, writing – review & editing, supervision, funding acquisition.

## Funding

J.G., A.S., and U.M. were supported by the John Templeton Foundation and the Fetzer Institute (Consciousness and Free Will: A Joint Neuroscientific-Philosophical Investigation (John Templeton Foundation #61283; Fetzer Institute, Fetzer Memorial Trust #4189)). J.G. was supported by the NSF (BCS-2219800).

## Declaration of Competing Interests

The authors have no competing interests to report.

## Supplementary Material

**Supplementary Figure 1:**
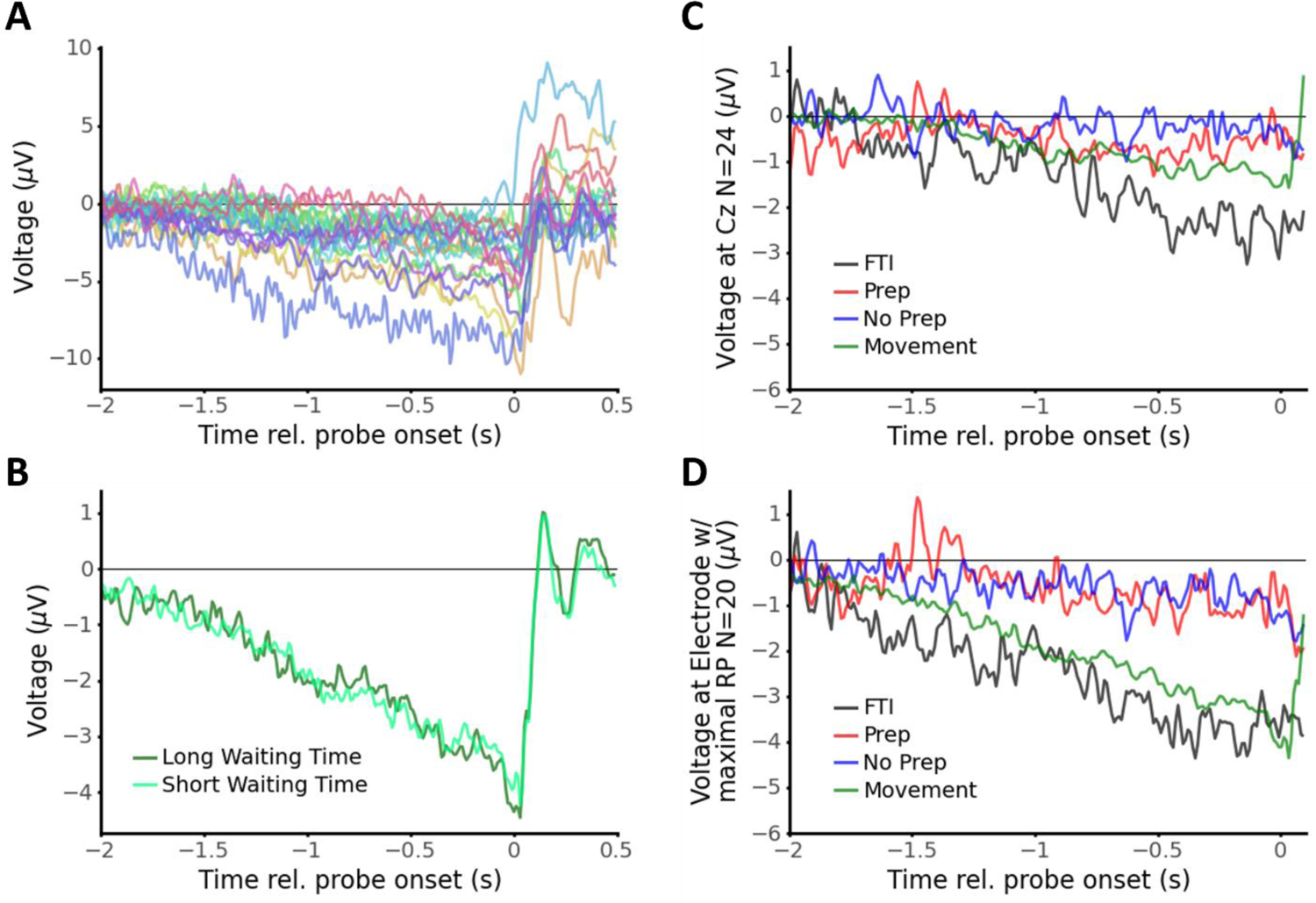
Supplementary ERP findings. A: Individual participants’ readiness potentials in Movement trials after selecting the electrode with maximal negativity (Cz n=9, FCz n=9, Fz n=2). B: Unlike Schurger (2018), we did not find a noticeable difference in RP amplitude for trials with relatively short or long waiting-times (by-participant median split after removing waiting times less than 2.5 seconds). Longer waiting times were more likely to be pre-empted by a probe, potentially leading to this lack of difference. C: Grand-averaged ERPs (averaged across trials, and then averaged across participants) for all 24 participants at electrode Cz. Lower RP amplitude emerges due to inclusion of participants that did not show an RP and selecting Cz instead of electrodes at which RP is maximal. D: Grand-averaged RP (averaged across trials, then averaged across participants using our selection criteria as in the main article) using traditional averaging methods instead of mixed-effects modeling. Findings are largely consistent with mixed-effects models.

**Supplementary Table 1:** Summary of estimated pre-probe EEG slope [−1s, 0s] and amplitudes [−0.25 s, 0s] (times relative to probe onset), each with their confidence intervals, for Prep, No Prep, and FTI conditions under different participant retention criteria. Also given are Bayes factors (BF_10_; Bayesian ANOVA in JASP), comparing RP amplitude in [−0.25s, 0s] across conditions. For all the retention criteria that we tested, when comparing Prep vs No Prep (bolded), we found Bayes factors around or below 0.1. This provides moderate to strong evidence that the EEG signals in the two conditions are the same, regardless of the retention criteria we used.

| Inclusion criteria | Condition | Slope<br>[-1s, 0s] | 95% CI | Amplitude<br>[-0.25s, 0s] | 95% CI | BF<sub>10</sub> vs<br>Prep | BF<sub>10</sub> vs No<br>Prep. | BF<sub>10</sub> vs<br>FTI |
| --- | --- | --- | --- | --- | --- | --- | --- | --- |
| Visible RP (n=20/24,<br>main text) | Prep. | -0.509<br>mV/s | (-0.904,<br>-0.115) | -1.102 mV | (-2.671,<br>0.466) | N/A | **0.091** | 1.584 |
|  | No Prep. | -0.661<br>mV/s | (-0.939,<br>-0.383) | -0.622 mV | (1.973,<br>0.729) | **0.091** | N/A | 21.008 |
|  | FTI | -1.994<br>mV/s | (-2.394,<br>-1.593) | -3.448 mV | (-5.019,<br>-1.877) | 1.584 | 21.008 | N/A |
| + At least 10 prep & no<br>prep trials each<br>(n=11/24) | Prep. | -0.582<br>mV/s | (-0.992,<br>-0.172) | -0.461 mV | (-2.087,<br>1.166) | N/A | **0.105** | 0.911 |
|  | No Prep. | -0.046<br>mV/s | (-0.395,<br>0.302) | -0.335 mV | (-1.868,<br>1.198) | **0.105** | N/A | 3.719 |
|  | FTI | -1.125<br>mV/s | (-1.716,<br>-0.535) | -3.549 mV | (-5.550,<br>-1.548) | 0.911 | 3.719 | N/A |
| + Frequency of being<br>probed between 30%<br>and 70% of trials<br>(n=12/24) | Prep. | -0.663<br>mV/s | (-1.136,<br>-0.191) | -0.430 mV | (-2.122,<br>1.262) | N/A | **0.108** | 11.706 |
|  | No Prep. | -0.327<br>mV/s | (-0.665,<br>0.010) | -0.958 mV | (-2.387,<br>0.470) | **0.108** | N/A | 14.823 |
|  | FTI | -2.054<br>mV/s | (-2.511,<br>-1.597) | -3.263 mV | (-4.929,<br>-1.597) | 11.706 | 14.823 | N/A |

**Supplementary Figure 2:**
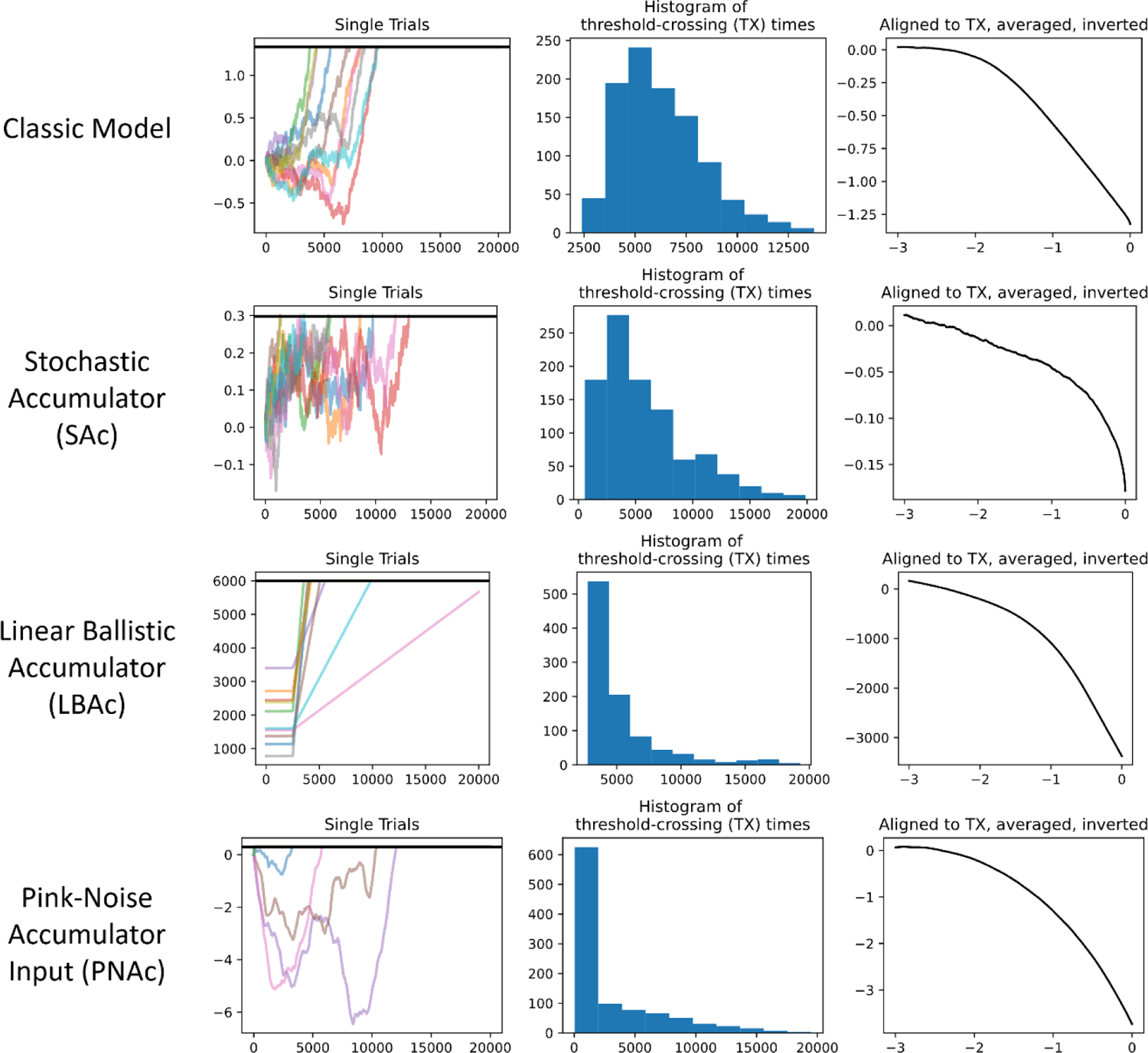
Some individual runs (left), the threshold-crossing distribution (middle), and threshold-aligned back-averages (right) of four different accumulator models. Classic model: noisy fluctuations occur until an ‘unconscious decision’ is made, after which a strong input drives activity towards the threshold. Stochastic accumulator (SAc): noisy fluctuations accumulate and trigger movement after crossing a threshold. Linear ballistic accumulator (LBAc): after 2500 ms, a linear drive (drawn from a normal distribution, in addition to a starting value) pushes activity and triggers movement after crossing a threshold. Pink-noise accumulator (PNAc): autocorrelated noise (1/f exponent = 1.5) accumulates and triggers movement when crossing a threshold. All four model types recreated a long-tail distribution of threshold-crossing times (or wait times), and all recreated an early deflection as seen in the RP. However, only SAc with dual-stage metacognition explained our EEG results (Fig. 4).

**Supplementary Figure 3:**
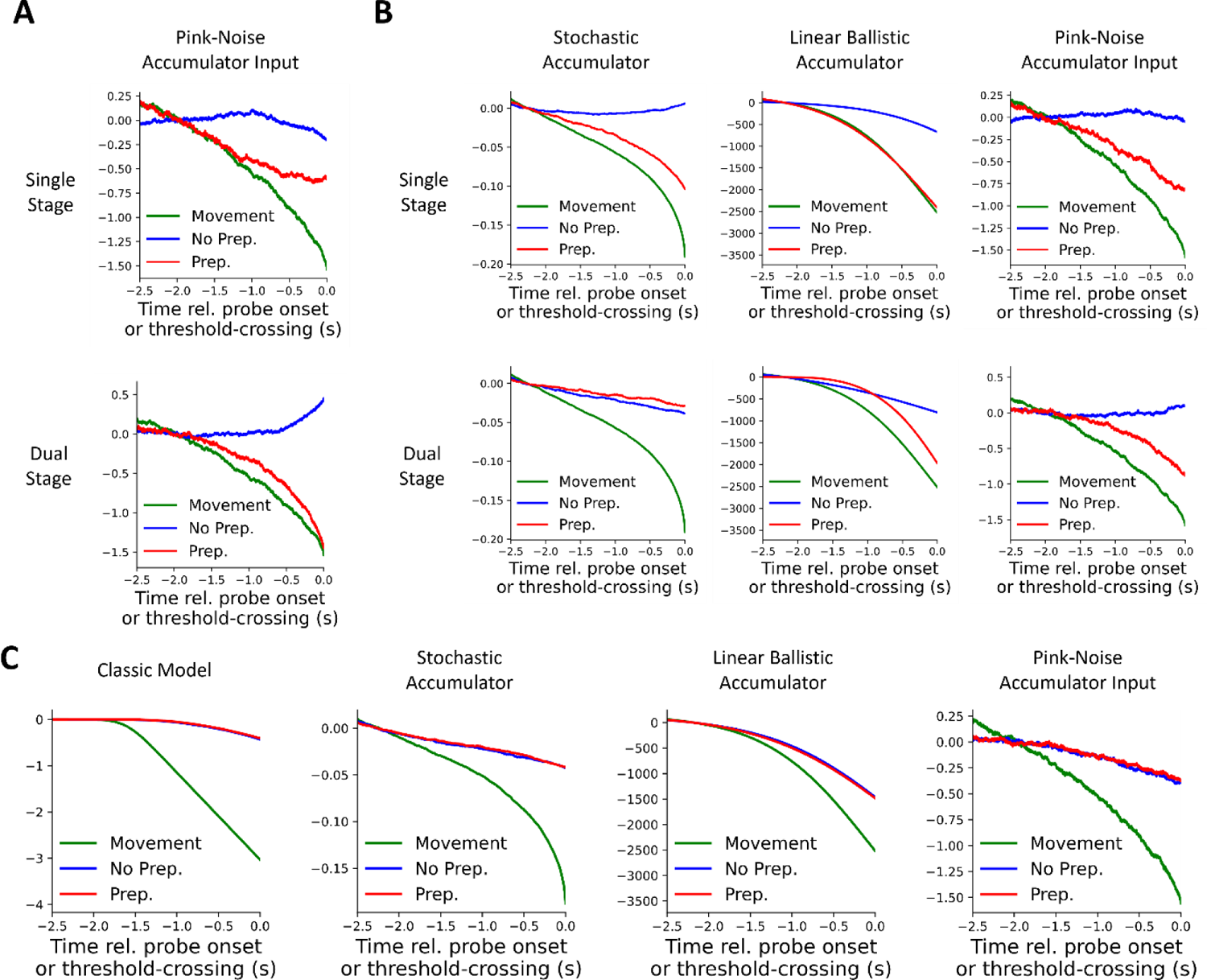
Supplementary modeling results. **A.** Pink-noise accumulator model^10^. In this model, the RP reflects an autocorrelated noisy input to a drift-diffusion process rather than the process itself. Aligned to threshold-crossings, the noisy input will also show an early negative deflection akin to the RP. As in Figure 4C, we show modeling results where the decision to report ‘Yes’ or ‘No’ in response to the probe is based on the amplitude (single-stage) or slope (dual-stage) of the accumulator (not on the noisy input) at or immediately after the time of the probe. Like our findings for the stochastic accumulator model, dual-stage metacognition with this process can explain our ERP results. Furthermore, we did not find a difference between RPs on trials where participants waited a short versus long amount of time to move (Fig. S1B), which was the crucial difference that used to support this model over the standard stochastic accumulator model^10^. **B.** Model results if the data were split according to accumulator amplitude (single-stage) or slope (dual-stage) 400 milliseconds following probe onset, rather than immediately following the probe. This would presumably be the case if some time was required for auditory processing and task-shifting. Though not identical, the results are largely the same with those obtained from the model without a delay (compare Fig. 4C). **C.** Model results if participants’ reports about awareness of preparation are based on unrelated brain activity *or* are pure guesses. We conducted similar analyses to Figure 4, except non-movement and non-FTI trials were randomly designated as Prep or No Prep, as one would expect if intention awareness was independent of the process that generates the RP or if reports were mere guesses. All four models were able to recreate our ERP results under these conditions (no difference between Prep and No Prep, and a negative slope before probe onset in both of those conditions). However, random guesses are unlikely because our participants reported being unsure on a relatively small percentage of trials (∼9%, Fig. 2). Furthermore, the probability of reporting that they were preparing to move increased with trial duration (p = 0.001; see behavioral results). We therefore think the most likely case is that participants metacognitively access stochastic accumulation following probe onset (Fig. 4) or base their reports on another feature of neural activity that is minimally dependent on the RP (e.g. beta power, Fig. 5). Nevertheless, these results clarify that a classic or linear ballistic model underlying the RP is mutually exclusive with reports of motor preparation being related to the RP at all—one cannot have it both ways.

## Notes

### Competing Interest Statement

The authors have declared no competing interest.

### Summary of Updates

We updated the manuscript structure, introduced new panels in several figures (including supplementary), conducted new analyses, and expanded our discussion.

## References

1. Kornhuber, H. H. & Deecke, L. [CHANGES IN THE BRAIN POTENTIAL IN VOLUNTARY MOVEMENTS AND PASSIVE MOVEMENTS IN MAN: READINESS POTENTIAL AND REAFFERENT POTENTIALS]. Pflugers Arch Gesamte Physiol Menschen Tiere 284, 1–17 (1965).

2. Libet, B., Gleason, C. A., Wright, E. W. & Pearl, D. K. Time of conscious intention to act in relation to onset of cerebral activity (readiness-potential). The unconscious initiation of a freely voluntary act. Brain 106 **(Pt** **3**), 623–642 (1983).

3. Sakata, H. et al. Slow Accumulations of Neural Activities in Multiple Cortical Regions Precede Self-Initiation of Movement: An Event-Related fMRI Study. eNeuro 4, ENEURO.0183-17.2017 (2017).

4. Soon, C. S., Brass, M., Heinze, H.-J. & Haynes, J.-D. Unconscious determinants of free decisions in the human brain. Nature Neuroscience 11, 543–545 (2008).

5. Haggard, P. & Eimer, M. On the relation between brain potentials and the awareness of voluntary movements. Exp Brain Res 126, 128–133 (1999).

6. Fried, I., Mukamel, R. & Kreiman, G. Internally Generated Preactivation of Single Neurons in Human Medial Frontal Cortex Predicts Volition. Neuron 69, 548–562 (2011).

7. Shibasaki, H. & Hallett, M. What is the Bereitschaftspotential? Clin Neurophysiol 117, 2341–2356 (2006).

8. Libet, B. Unconscious cerebral initiative and the role of conscious will in voluntary action. THE BEHAVIORAL AND BRAIN SCIENCES 38 (1985).

9. Schurger, A., Sitt, J. D. & Dehaene, S. An accumulator model for spontaneous neural activity prior to self-initiated movement. PNAS 109, E2904–E2913 (2012).

10. Schurger, A. Specific Relationship between the Shape of the Readiness Potential, Subjective Decision Time, and Waiting Time Predicted by an Accumulator Model with Temporally Autocorrelated Input Noise. eNeuro ENEURO.0302-17.2018 (2018) doi:10.1523/ENEURO.0302-17.2018.

11. Khalighinejad, N., Schurger, A., Desantis, A., Zmigrod, L. & Haggard, P. Precursor processes of human self-initiated action. NeuroImage 165, 35–47 (2018).

12. Maoz, U., Yaffe, G., Koch, C. & Mudrik, L. Neural precursors of decisions that matter—an ERP study of deliberate and arbitrary choice. eLife 8, e39787 (2019).

13. Moutard, C., Dehaene, S. & Malach, R. Spontaneous Fluctuations and Non-linear Ignitions: Two Dynamic Faces of Cortical Recurrent Loops. Neuron 88, 194–206 (2015).

14. Schmidt, S., Jo, H.-G., Wittmann, M. & Hinterberger, T. ‘Catching the waves’ – slow cortical potentials as moderator of voluntary action. Neuroscience & Biobehavioral Reviews 68, 639–650 (2016).

15. Schurger, A., Hu, P. ‘Ben’, Pak, J. & Roskies, A. L. What Is the Readiness Potential? Trends in Cognitive Sciences 25, 558–570 (2021).

16. Gavenas, J., Rutishauser, U., Schurger, A. & Maoz, U. Slow ramping emerges from spontaneous fluctuations in spiking neural networks. 2023.05.27.542589 Preprint at 10.1101/2023.05.27.542589 (2023).

17. Bogler, C., Grujičić, B. & Haynes, J.-D. Confusions regarding stochastic fluctuations and accumulators in spontaneous movements. 2021.06.04.447111 Preprint at 10.1101/2021.06.04.447111 (2021).

18. Brown, S. D. & Heathcote, A. The simplest complete model of choice response time: linear ballistic accumulation. Cogn Psychol 57, 153–178 (2008).

19. Dominik, T., Mele, A., Schurger, A. & Maoz, U. Libet’s legacy: A primer to the neuroscience of volition. Neuroscience & Biobehavioral Reviews 157, 105503 (2024).

20. Dominik, T. et al. Libet’s experiment: Questioning the validity of measuring the urge to move. Consciousness and Cognition 49, 255–263 (2017).

21. Schlegel, A. et al. Barking up the wrong free: readiness potentials reflect processes independent of conscious will. Exp Brain Res 229, 329–335 (2013).

22. Bredikhin, D., Germanova, K., Nikulin, V. & Klucharev, V. (Non)-experiencing the intention to move: On the comparisons between the Readiness Potential onset and Libet’s W-time. Neuropsychologia 108570 (2023) doi:10.1016/j.neuropsychologia.2023.108570.

23. Parés-Pujolràs, E., Kim, Y.-W., Im, C.-H. & Haggard, P. Latent awareness: Early conscious access to motor preparation processes is linked to the readiness potential. NeuroImage 202, 116140 (2019).

24. Schultze-Kraft, M., Parés-Pujolràs, E., Matić, K., Haggard, P. & Haynes, J.-D. Preparation and execution of voluntary action both contribute to awareness of intention. Proceedings of the Royal Society B: Biological Sciences 287, 20192928 (2020).

25. Parés-Pujolràs, E., Matić, K. & Haggard, P. Feeling ready: neural bases of prospective motor readiness judgements. Neuroscience of Consciousness 2023, niad003 (2023).

26. Matsuhashi, M. & Hallett, M. The timing of the conscious intention to move. Eur J Neurosci 28, 2344–2351 (2008).

27. Verbaarschot, C., Haselager, P. & Farquhar, J. Detecting traces of consciousness in the process of intending to act. Exp Brain Res 234, 1945–1956 (2016).

28. Triggiani, A. I. et al. What is the Intention to Move and When Does it Occur? Neuroscience & Biobehavioral Reviews 105199 (2023) doi:10.1016/j.neubiorev.2023.105199.

29. Banks, W. P. & Isham, E. A. We infer rather than perceive the moment we decided to act. Psychol Sci 20, 17–21 (2009).

30. Lau, H. C., Rogers, R. D. & Passingham, R. E. Manipulating the Experienced Onset of Intention after Action Execution. Journal of Cognitive Neuroscience 19, 81–90 (2007).

31. Maoz, U. et al. On reporting the onset of the intention to move. Surrounding free will: philosophy, psychology, neuroscience (Mele A, ed) 184–202 (2015).

32. Pfurtscheller, G. & Aranibar, A. Event-related cortical desynchronization detected by power measurements of scalp EEG. Electroencephalogr Clin Neurophysiol 42, 817– 826 (1977).

33. Pfurtscheller, G. & Lopes da Silva, F. H. Event-related EEG/MEG synchronization and desynchronization: basic principles. Clin Neurophysiol 110, 1842–1857 (1999).

34. Park, H.-D. et al. Breathing is coupled with voluntary action and the cortical readiness potential. Nature Communications 11, 1–8 (2020).

35. Garipelli, G., Chavarriaga, R. & Millán, J. del R. Single trial analysis of slow cortical potentials: a study on anticipation related potentials. J. Neural Eng. 10, 036014 (2013).

36. Mike X Cohen. Analyzing Neural Time Series Data: Theory and Practice. (The MIT Press, 2014).

37. Schultze-Kraft, M. et al. The point of no return in vetoing self-initiated movements. PNAS 113, 1080–1085 (2016).

38. JASP Team (2023). JASP.

39. Moran, R., Teodorescu, A. R. & Usher, M. Post choice information integration as a causal determinant of confidence: Novel data and a computational account. Cognitive Psychology 78, 99–147 (2015).

40. Desender, K., Ridderinkhof, K. R. & Murphy, P. R. Understanding neural signals of post-decisional performance monitoring: An integrative review. eLife 10, e67556 (2021).

41. Pleskac, T. J. & Busemeyer, J. R. Two-stage dynamic signal detection: A theory of choice, decision time, and confidence. Psychological Review 117, 864–901 (2010).

42. Jolly, E. Pymer4: Connecting R and Python for Linear Mixed Modeling. JOSS 3, 862 (2018).

43. Bai, O. et al. Prediction of human voluntary movement before it occurs. Clinical Neurophysiology 122, 364–372 (2011).

44. Lew, E., Chavarriaga, R., Zhang, H., Seeck, M. & Millán, J. del R. Self-paced movement intention detection from human brain signals: Invasive and non-invasive EEG. in 2012 Annual International Conference of the IEEE Engineering in Medicine and Biology Society 3280–3283 (2012). doi:10.1109/EMBC.2012.6346665.

45. Lew, E. Y. L., Chavarriaga, R., Silvoni, S. & Millán, J. del R. Single trial prediction of self-paced reaching directions from EEG signals. Frontiers in Neuroscience 8, (2014).

46. Shenoy, K. V., Kaufman, M. T., Sahani, M. & Churchland, M. M. A dynamical systems view of motor preparation: Implications for neural prosthetic system design. Prog Brain Res 192, 33–58 (2011).

47. Neafsey, E. J. Conscious intention and human action: Review of the rise and fall of the readiness potential and Libet’s clock. Consciousness and Cognition 94, 103171 (2021).

48. Verbaarschot, C., Haselager, P. & Farquhar, J. Probing for Intentions: Why Clocks Do Not Provide the Only Measurement of Time. Frontiers in Human Neuroscience 13, (2019).

49. Verbruggen, F. & Logan, G. D. Response inhibition in the stop-signal paradigm. Trends Cogn Sci 12, 418–424 (2008).

50. Mosher, C. P., Mamelak, A. N., Malekmohammadi, M., Pouratian, N. & Rutishauser, U. Distinct roles of dorsal and ventral subthalamic neurons in action selection and cancellation. Neuron 109, 869–881.e6 (2021).

51. Raud, L. et al. A Single Mechanism for Global and Selective Response Inhibition under the Influence of Motor Preparation. J. Neurosci. 40, 7921–7935 (2020).

52. Wagner, J., Wessel, J. R., Ghahremani, A. & Aron, A. R. Establishing a Right Frontal Beta Signature for Stopping Action in Scalp EEG: Implications for Testing Inhibitory Control in Other Task Contexts. Journal of Cognitive Neuroscience 30, 107–118 (2018).

53. Wessel, J. R. β-Bursts Reveal the Trial-to-Trial Dynamics of Movement Initiation and Cancellation. J. Neurosci. 40, 411–423 (2020).

54. Wokke, M. E., Cleeremans, A. & Ridderinkhof, K. R. Sure I’m Sure: Prefrontal Oscillations Support Metacognitive Monitoring of Decision Making. J. Neurosci. 37, 781–789 (2017).

55. Soutschek, A., Moisa, M., Ruff, C. C. & Tobler, P. N. Frontopolar theta oscillations link metacognition with prospective decision making. Nat Commun 12, 3943 (2021).

56. Tang, S. et al. The dynamic monitoring and control mechanism in problem solving: Evidence from theta and alpha oscillations. International Journal of Psychophysiology 170, 112–120 (2021).

57. Wokke, M. E., Achoui, D. & Cleeremans, A. Action information contributes to metacognitive decision-making. Sci Rep 10, 3632 (2020).

58. Brus, J., Aebersold, H., Grueschow, M. & Polania, R. Sources of confidence in value-based choice. Nat Commun 12, 7337 (2021).

59. Dippel, G., Mückschel, M., Ziemssen, T. & Beste, C. Demands on response inhibition processes determine modulations of theta band activity in superior frontal areas and correlations with pupillometry – Implications for the norepinephrine system during inhibitory control. NeuroImage 157, 575–585 (2017).

60. Pertermann, M., Mückschel, M., Adelhöfer, N., Ziemssen, T. & Beste, C. On the interrelation of 1/f neural noise and norepinephrine system activity during motor response inhibition. Journal of Neurophysiology 121, 1633–1643 (2019).

61. Cavanagh, J. F. & Frank, M. J. Frontal theta as a mechanism for cognitive control. Trends in Cognitive Sciences 18, 414–421 (2014).

